# Resistance of estrogen receptor function to BET bromodomain inhibition is mediated by transcriptional coactivator cooperativity

**DOI:** 10.1101/2024.07.25.605008

**Authors:** Sicong Zhang, Robert G. Roeder

## Abstract

The Bromodomain and Extra-Terminal Domain (BET) family of proteins are critical chromatin readers that bind to acetylated histones through their bromodomains to activate transcription. Here, we reveal that bromodomain inhibition fails to repress oncogenic targets of estrogen receptor due to an intrinsic transcriptional mechanism. While bromodomains are necessary for the transcription of many genes, BRD4 binds to estrogen receptor binding sites and activates transcription of critical oncogenes independently of its bromodomains. BRD4 associates with the Mediator complex and disruption of Mediator complex reduces BRD4’s enhancer occupancy. Profiling changes in the post-initiation RNA polymerase II (Pol II)-associated factors revealed that BET proteins regulate interactions between Pol II and elongation factors SPT5, SPT6, and PAF1 complex, which associate with BET proteins independently of their bromodomains and mediate their transcription elongation effect. Our findings highlight the importance of bromodomain-independent functions and interactions of BET proteins in the development of future therapeutic strategies.

## Introduction

Activator-dependent transcription relies on specific co-activators, and targeting these co-activators pharmacologically offers a promising therapeutic approach to modulate transcriptional programs and selectively attenuate disease-causing gene expression^1^. One prominent group of transcriptional co-activators is the BET family of proteins, comprising four members: BRD2, BRD3, BRD4, and the testis-specific BRDT. BET proteins are known as key readers of chromatin acetylation. They possess two tandem bromodomains, BD1 and BD2, that bind to acetyl-lysine residues, and an Extra-Terminal (ET) domain that interacts with various transcription regulators^2,3^. The bromodomains of BET proteins play two major roles in transcriptional regulation. First, recruitment of BET proteins to chromatin is generally presumed to occur through bromodomain interactions with acetylated histones that are enriched at enhancers and promoters of active genes^4^. Second, bromodomains are essential for BET proteins to facilitate transcription through an intrinsic histone-chaperone activity^5,6^. Apart from bromodomain interactions, the ET domain of BET proteins bind multiple transcriptional regulators^2,7^ that function as “effectors” to regulate transcription after BET proteins are recruited to chromatin. BRD4 has been extensively studied because of its crucial role in supporting *MYC* transcription^8^. It is known to bind acetylated histones to activate transcriptional elongation by recruiting positive transcription elongation factor b (P-TEFb) to chromatin through its C-terminal end^9^. However, recent findings have suggested that BET proteins act as master transcription elongation factors independent of CDK9 recruitment^10^, such that the exact mechanisms of how BET proteins regulate elongation are not fully understood. BET inhibitors (BETi) competitively bind to bromodomains, displacing BET proteins from chromatin and disrupting oncogene transcription. Bromodomain inhibition by BETi has demonstrated potent antitumor activities in various cancers by suppressing expression of oncogenes, such as *MYC* in leukemia^8,11^. However, recent clinical trials involving estrogen receptor-positive (ER+)/HER2-breast cancer patients have shown that BETi do not exhibit efficacy (ref ^12^ and ClinicalTrials.gov/NCT02964507). In ER+ breast cancer, ER plays a pivotal role in promoting tumor growth by activating the transcription of critical oncogenes that include *MYC* and *CCND1*^13^. Gene expression profiles of BETi-treated tumors from ER+ breast cancer patients have revealed the persistence or activation of a proliferation program controlled by MYC and E2F, despite the effective suppression of various inflammatory genes that are well-known targets of BETi^12,14^. Interestingly, BETi have been reported to inhibit ER-mediated transcription of certain genes^15–17^, but conflicting results based on knockdown experiments have been reported regarding which BET protein is required for ER-mediated transcription -- with one study showing an essential role of BRD4 and another implicating BRD3 as a major player^16,17^. These observations underscore the imperative need for a comprehensive investigation into the molecular mechanisms governing BET proteins in ER-mediated transcription. This endeavor not only stands to provide a model for transcriptional studies but also holds significant promise for advancing future drug development efforts.

Here, we unveil that the ER-mediated transcription of numerous genes, including *MYC*, exhibits intrinsic resistance to bromodomain inhibition due to the bromodomain-independent recruitment and activity of BET proteins through other transcriptional co-activators. Our findings shed light on the complex regulatory mechanisms of ER-mediated transcription and offer explanations for the limited efficacy of BETi in ER+ breast cancer patients. Understanding these mechanisms can inform novel strategies to overcome endocrine resistance and improve therapeutic outcomes in various cancer patients, including breast cancer.

## Results

### Transcription of a large group of ER target genes is resistant to BETi

Considering the lack of efficacy of BETi, it is likely that BETi fail to fully repress the ER-mediated transcription of critical oncogenic targets through an intrinsic mechanism or that these genes can be activated by other transcription factors that are resistant to BETi through an adaptive mechanism. To distinguish these possibilities, we re-analyzed the gene expression profile in MCF7 cells, a human ER+ breast cancer cell line, treated with the pan-BETi JQ1 for short time from existing datasets^16,17^. Despite different treatment conditions in two independent studies, both studies revealed that while some ER-regulated genes were indeed repressed by BETi, many important oncogenic targets such as *MYC* and *CCND1* remained highly induced by estrogen (Fig. 1a and Extended Data Fig. 1a). Moreover, excluding genes that were upregulated by JQ1 alone, 35-44% of E_2_-upregulated genes showed BETi resistance, defined by higher expression in combination with JQ1 and estrogen relative to control; these included 42 common genes, such as *MYC* and *RET,* in both studies (Fig. 1b). These analyses suggest that the failure of BETi is likely caused by an intrinsic resistance at the beginning of BETi treatment. Two possible mechanisms can be envisioned to explain the BETi resistance. It could result from a dispensability either of BET proteins or of bromodomains in ER-dependent transcription of a group of genes – even though these genes, such as *MYC,* require BET proteins in other cancers.

**Fig. 1.**
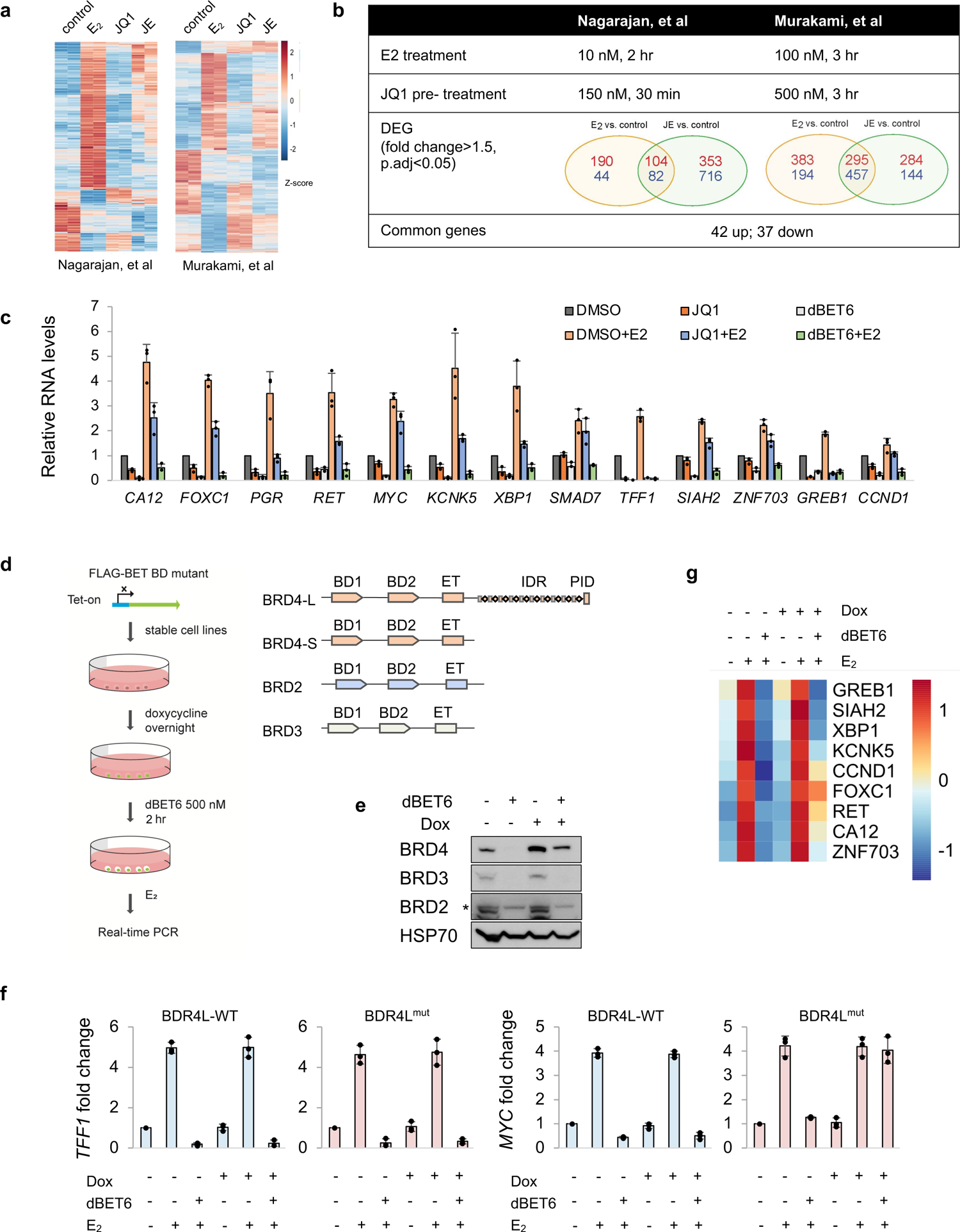
ER-mediated transcriptional resistance to bromodomain inhibition. **a**, Heatmap showing effect of JQ1 on ER target gene expression analyzed from two studies under conditions described in **b**. **c**, qPCR of ER target genes in MCF7 cells using primers for pre-mRNA (except for *MYC,* whose pre-mRNA and mRNA levels show identical change, and *FOXC1*, which is a single-exon gene.) **d**, Schematic of Tet-on system for expression of single BET BD mutants. Doxycycline was added at 100 ng/ml overnight to induce transgene expression in MCF7/FRT cells. Schematic of BRD4 indicating the two bromodomains (BD1 and BD2), the Extra-Terminal domain (ET), the intrinsic disordered region (IDR) and the P-TEFb-interacting domain (PID). **e**, Immunoblot of BET proteins in MCF7/FRT-BRD4L^mut^ cells treated with doxycycline and dBET6. **f**, qPCR of gene expression changes in MCF7/FRT-BRD4L-WT and MCF7/FRT-BRD4L^mut^ cells. **g**, Heatmap showing effect of BRD4L^mut^ expression on ER-mediated gene transcription measured by qPCR. Whereas BRD4L^mut^ showed no rescue of *GREB1* transcription and minimal rescue of *SIAH2* and *XBP1* transcription, it partially rescued other gene transcription and fully restored *FOXC1* transcription. For all RNA analyses by qPCR, cells were pre-treated with 500 nM JQ1, dBET6, or dTAG-7 (or dTAG^V^-1) for 2 hours and then stimulated with 10 nM estradiol (E_2_) for 1 hour except where otherwise stated. Data shown as mean ± s.d. from independent triplicates. See also Extended Data Fig.1.

To compare the requirement of BET proteins vs. their bromodomains, we measured the effects of JQ1 and pan-BET degraders on ER-mediated transcription of select BETi-sensitive and -resistant genes by quantitative reverse transcription-PCR (qRT-PCR). The pan-BET degraders MZ1 and dBET6^10,18^ rapidly depleted BET proteins without affecting ER protein levels (Extended Data Fig.1b,c). For the BETi-sensitive genes *TFF1* and *GREB1,* both JQ1 and pan-BET degraders completely blocked the E_2_-induced transcription. For the BETi-resistant genes *MYC* and *FOXC1*, dBET6 and MZ1 were far more potent than JQ1 in reducing E_2_-induced transcription, indicating an indispensable role of BET proteins (Fig. 1c and Extended Data Fig. 1d). It therefore is likely that bromodomain-independent mechanisms, rather than the dispensability of the BET proteins themselves, account for the chromatin binding and function of BET proteins in JQ1 treatment.

### BRD4L enables BETi-resistant ER-mediated gene transcription

Next, we set out to determine which BET protein is responsible for the ER-mediated transcription of BETi-resistant genes. In contrast to early studies^16,17^, we found that shRNA-mediated knockdowns of individual BET proteins did not affect the levels of other BET proteins, ER or H2Bub1, a histone mark of transcription elongation (Extended Data Fig.1e-g). To circumvent the confounding effects from long-term experiments, we devised a strategy (Fig. 1d) involving transient expression of a BET bromodomain mutant^19^ along with the use of the bromodomain-dependent BET degrader to deplete all endogenous BET proteins. This approach allowed us to directly evaluate the function of individual BET proteins. We first derived MCF7/FRT/Tet-on cell lines expressing either a FLAG-tagged wild type (WT) BRD4 or a bromodomain mutant (BD) BRD4 under the control of doxycycline. dBET6 depleted all endogenous BET proteins and WT BRD4 without affecting the BD mutant (Fig. 1e and Extended Data Fig. 1h). Encouraged by this result, we further generated cell lines expressing bromodomain mutants of BRD2 and BRD3, which were validated as being resistant to dBET6 as well. The BD mutant of BRD4 long isoform (BRD4L^mut^), but not WT BRD4 (BRD4L-WT) (Fig. 1f) or BD mutants of other BET proteins (data not shown), efficiently rescued the loss of ER-mediated expression of *MYC* in the presence of dBET6. Although BRD4L^mut^ failed to restore *TFF1* and *GREB1* transcription, it rescued many BETi-resistant genes to various extents (Fig. 1f,g). These experiments provide direct evidence in support of the ability of BRD4 to assist ER-mediated transcription in a bromodomain-independent manner, which gives rise to the intrinsic BETi resistance of some genes. Of note, expression of these BD mutants under the control of Tet-On 3G transactivator substantially repressed ER-mediated transcription likely due to the squelching effect^20^ (Extended Data Fig.1i). As Tet-on systems are widely used in transcriptional studies, careful analysis and interpretation are necessary for transactivator-based assays.

Next, we used the degradation tag (dTAG) strategy^21^ to ask if rapid depletion of endogenous BRD4 could reverse the BETi-resistance of these genes (Extended Data Fig.2a,b). dTAG compounds (dTAG7 and dTAG^V^-1) targeting BRD4 efficiently and selectively depleted BRD4 without significantly decreasing global levels of H2Bub1 or the elongating (CTD Ser2-P hyperphosphorylated) form of Pol II (Extended Data Fig.2c,d). Rapid depletion of BRD4 reduced ER-mediated transcription of *FOXC1* and *MYC* to levels comparable to those following JQ1 treatment, and reduced *TFF1* to a much less extent than JQ1 (Fig. 2a). When combined with JQ1, BRD4 depletion further reduced transcription to levels comparable to those observed with dBET6 treatment, suggesting redundant roles of BET proteins in ER-mediated transcription and the selective requirement for bromodomains by different BET proteins at different ER target genes (Fig. 2a). These observations imply that for BETi-sensitive genes, for which bromodomains are absolutely required, BRD4 is recruited or functions along with other BET proteins through bromodomains. For BETi-resistant genes, such as *MYC* and *FOXC1*, BRD4L can function independently of its bromodomains, whereas the function of other BET proteins must rely on bromodomains.

**Fig. 2.**
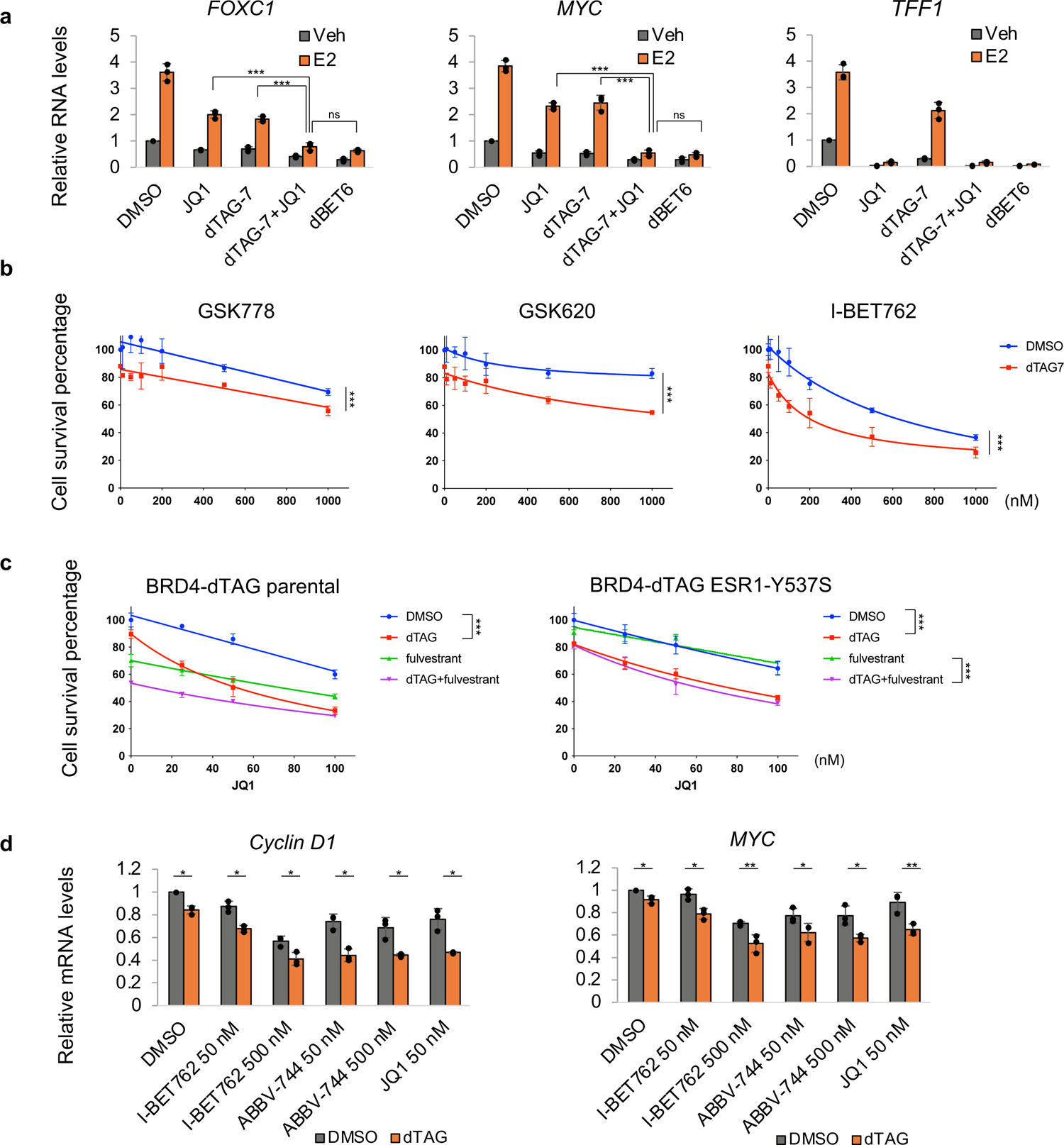
BRD4 depletion potentiates BETi response. **a**, qPCR of gene expression changes in BRD4-dTAG cells. **b**, Percentage of MCF7 BRD4-dTAG cell survival after treatment with or without BET inhibitors and dTAG-7 (250 nM) for 6 days. **c**, Percentage of BRD4-dTAG parental or ESR1-Y537S expressing cell survival after treatment with JQ1, fulvestrant (0.5 nM) or dTAG-7 (250 nM) for 6 days. **d**, qPCR of *MYC* and *Cyclin D1* mRNA levels in MCF7 BRD4-dTAG ESR1-Y537S cells upon overnight treatment with BET inhibitors and dTAG-7 (500 nM). Cells in **b-d** were grown in normal full medium containing 10% FBS. Data shown as mean ± s.d. from independent triplicates. p values from two-tailed t-tests are depicted with asterisks. See also Extended Data Fig.2.

### BRD4 depletion potentiates BETi response

To investigate the potential enhanced antitumor activity of BETi treatment in conjunction with BRD4 depletion, we conducted a comparative analysis of the anti-proliferative effects of various BETi, both individually and in combination with BRD4 depletion. When BRD4 depletion alone was tested, it resulted in a modest 10% reduction in cell growth (Fig. 2b). However, this depletion significantly augmented the anti-proliferation activity of all tested BETi in MCF7 cells. Our findings also indicated that BD2-preferred ABBV-744 and pan-BD inhibitors I-BET762 and JQ1 exhibited greater potency than BD1-selective GSK778 and BD2-selective GSK620 inhibitors^22,23^ (Fig. 2b and Extended Data Fig. 2e). Furthermore, long-term exposure to JQ1 suppressed ER expression^15^, a phenomenon further accentuated by BRD4 depletion (Extended Data Fig.2f). In order to differentiate the impact on ER-mediated transcription from the alteration in ER expression, we introduced a transgene encoding the *ESR1* Y537S mutant, a constitutively active ER mutant found in endocrine-resistant tumors^24^. Fulvestrant, a selective estrogen receptor degrader, synergistically enhanced the anti-growth effect of BRD4 depletion and JQ1 in parental cells (Fig. 2c). However, fulvestrant displayed no effect in cells with the *ESR1* Y537S mutant, which is resistant to degradation by fulvestrant, suggesting that the effects observed with BETi and BRD4 depletion were not affected by ER expression in these cells (Fig. 2c). Consistent with the observed anti-proliferation effects, the combination of BRD4 depletion with I-BET762, ABBV-744, or JQ1 exhibited a more potent inhibition of *MYC* and *CCND1* expression compared to monotherapy in cells with the *ESR1* Y537S mutation, with a dose-dependent response observed for I-BET762 (50-500 nM) but not ABBV-744 (Fig. 2d and Extended Data Fig. 2g). The combinatorial inhibition of *MYC* and *CCND1* expression by BRD4 depletion and BETi were also observed for GSK778 and GSK620 (Extended Data Fig.2h). We also found a dose-dependent (0.1-1 µM) inhibition of *MYC* and *CCND1* expression by GSK778. In contrast, GSK620 showed the same inhibitory effects at 0.1 µM and 1 µM, and less potency compared to GSK778 at 1 µM, suggesting that inhibition of *MYC* and *CCND1* expression by pan-BETi was primarily attributed to BD1 inhibition (Extended Data Fig.2h). Collectively, our findings suggest that BRD4 depletion sensitizes tumor cells to the response of BETi, thereby potentially allowing a reduction in the therapeutic dosage of BETi and leading to improved safety and efficacy.

### BRD4L is recruited to ER binding sites

To understand the mechanism of bromodomain independency, we conducted ChIP-seq analyses to assess the recruitment of BRD4 to ER target genes using a commonly used antibody that recognizes the BRD4L C-terminus. We identified ER binding sites (ERBSs) from two existing ER ChIP-seq datasets that utilized different treatment conditions, antibodies, and peak calling algorithms. These datasets yielded both shared regions (representing core ERBSs) and non-overlapping regions (representing dynamic ERBSs). Among the 6,985 core ERBSs, we categorized them into high-ERBSs and low-ERBSs based on ER binding signals. In the absence of E_2_ stimulation, BRD4L exhibited minimal and comparable enrichment at both low-ERBSs and high-ERBSs (Fig. 3a). However, upon ligand stimulation, BRD4L binding to ERBSs significantly increased, with a greater enhancement observed at high-ERBSs compared to low-ERBSs (Fig. 3a). Treatment with JQ1 alone abolished the basal BRD4L binding, while E_2_ stimulation led to a notable increase in BRD4 binding at high-ERBSs and, to a lesser extent, at low-ERBSs (Fig. 3b). These observations were consistent with data on well-characterized ER-target genes, such as *TFF1* and *FOXC1* (Fig. 3c). *TFF1* contains three ERBSs, including a major distal enhancer (ERE3) located approximately 10 kb upstream of the transcription start site (TSS) and a promoter-proximal enhancer (TSS −400 bp). *FOXC1*, on the other hand, is a single-exon gene regulated by a distal enhancer. At these enhancers, treatment with JQ1 reduced but, particularly at the *FOXC1* enhancer, did not completely eliminate ER-dependent BRD4 binding. Next, we induced BRD4L^mut^ expression in MCF7/FRT/Tet-on cells by overnight doxycycline treatment and then conducted an N-terminal FLAG ChIP-seq analysis. The ChIP-seq peaks of BRD4L^mut^ overlapped with 72% of the endogenous BRD4 binding sites identified in triplicates (Fig. 3d). Moreover, the non-overlapping regions displayed significantly lower signal intensity compared to the overlapping sites (Fig. 3e). Hence, it can be concluded that the bromodomain-deficient BRD4 can occupy a majority of the BRD4 binding sites. At ERBSs, estrogen stimulation resulted in increased binding of BRD4L^mut^ (Fig. 3f,g), suggesting that BRD4L^mut^ is still selectively recruited to specific chromatin sites to play a role in transcription rather than acting through random encounters with the transcription machinery. Since the cells expressing BRD4L^mut^ also expressed endogenous BRD4, it was presumed that wild-type BRD4 would exhibit higher binding than BRD4L^mut^ and, consequently, that depletion of wild-type BRD4 would enhance the binding of BRD4L^mut^. Surprisingly, however, treatment with dBET6 led to an overall reduction in BRD4L^mut^ binding, except at *MYC* enhancers that showed less susceptibility to the effects of dBET6. These results indicate that wild-type BRD4 may facilitate the binding of BRD4L^mut^ and that bromodomains play a role in BRD4 self-association.

**Fig. 3.**
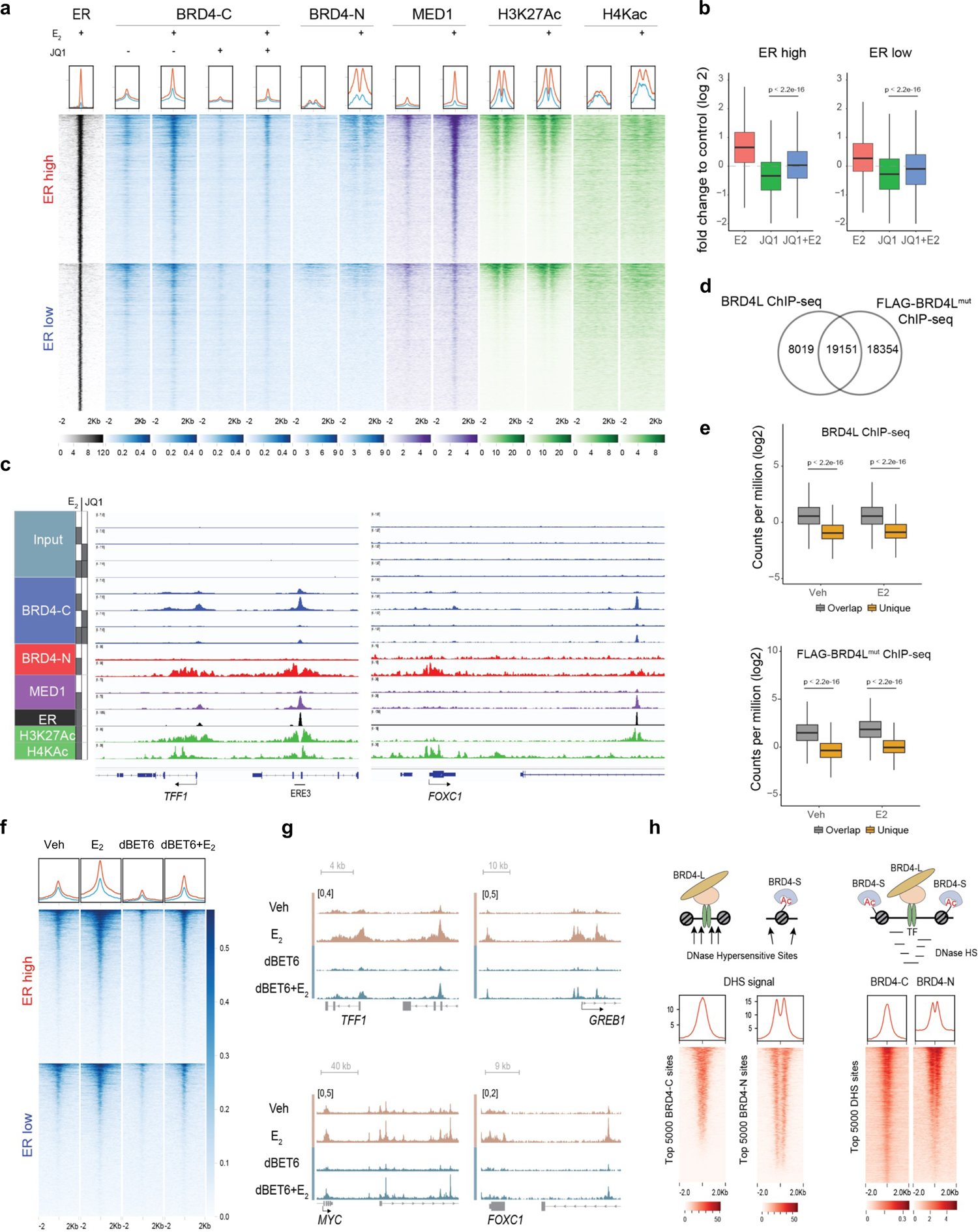
BRD4L is recruited to ERBSs. **a**, Heatmap of ER (black), BRD4 (blue), MED1 (purple) and Ac-H3/H4 (green) levels at ERBSs. **b**, Boxplot quantification of BRD4 ChIP-seq signals at ERBSs. p value from Welch Two Sample t-test. **c**, ChIP-seq signals of ER, BRD4-C (blue), BRD4-N (red), MED1 and Ac-H3/H4 at *FOXC1* and *TFF1* loci. ERE3 is the major enhancer of *TFF1*. **d**, Venn diagram showing overlapping binding sites from ChIP-seq of BRD4L and FLAG-BRD4L^mut^. **e**, Normalized tag counts at BRD4L and FLAG-BRD4L^mut^ overlapping or non-overlapping regions. **f**, Heatmap of FLAG-BRD4L^mut^ at ERBSs under indicated conditions. **g**, ChIP-seq signals of FLAG-BRD4L^mut^ at select gene loci. **h**, Left: heatmap of DNase hypersensitivity signal (DHS) at BRD4 binding sites from BRD4-N or BRD4-C ChIP-seq. Right: heatmap of BRD4-N and BRD4-C ChIP-seq signals at DNase hypersensitivity sites. For all ChIP performed in this study, cells were pre-treated with 500 nM of different compounds (e.g., JQ1, dBET6) for 2 hours and then stimulated with 10 nM E_2_ for 40 min except where otherwise stated. See also Extended Data Fig.3.

Our BRD4L ChIP-seq results yielded unexpected findings that contradicted results of a previous report of sites of BRD4 enrichment^16^. In this regard, and surprisingly, we observed co-localization of BRD4L (BRD4-C) with ER and Mediator subunit MED1 (Fig. 3a), whereas the other study^16^ indicated a connection between BRD4 and acetylated histones surrounding ERBSs (as observed for BRD4-N in Fig. 3a). For instance, at the *FOXC1* locus, the BRD4-C and BRD4-N ChIP-seq results exhibited nearly exclusive outcomes (Fig. 3c, blue vs. red). Our study thus identifies enhancer enrichment of BRD4L, while the other study detected binding regions that largely overlapped with acetylated histone regions that flanked ERBSs and extended from the promoter into the gene body. These conflicting outcomes can be attributed to the use of different antibodies. In our study, we utilized a commercial antibody that is specific to the C-terminal region of BRD4 (aa 1312–1362) and exclusively targets BRD4L. In contrast, the other study employed an N-terminal antibody (aa 149–284) that recognizes both isoforms (BRD4L and BRD4S). The divergent patterns generated by these two antibodies suggest that the N-terminal antibody fails to recognize BRD4 at ERBSs, while the C-terminal antibody either fails to detect BRD4L at ERBS-flanking histones and gene bodies or indicates a genuinely low enrichment of BRD4L in those regions compared to ERBSs. Upon closer examination of gene bodies of *TFF1, FOXC1*, and *MYC*, we observed higher signals of BRD4L and BRD4L^mut^ relative to nearby untranscribed regions but substantially lower signals compared to ERBSs. Hence, BRD4L exhibits greater enrichment at ERBSs in comparison to ERBS-flanking histones and gene bodies. This limited accessibility of certain antibodies to the N-terminal region of BRD4L may be attributed to its unique conformation or its distinct binding profile with certain factors.

Next, we investigated whether the enhancer enrichment of BRD4L extends to other transcription factors. To explore this question, we examined the relationship between chromatin accessibility and BRD4 binding in MCF7 cells and, intriguingly, found a significant association— with DNase I hypersensitivity (DHS) peaks being consistently centered over BRD4L binding sites, and vice versa (Fig. 3h). Similarly, when excluding ERBSs, BRD4L exhibited binding at the center of the ATAC (assay for transposase-accessible chromatin) regions (Extended Data Fig.3a). Furthermore, in MDA-MB-231 cells, an ER-negative breast cancer cell line, BRD4L exhibited binding at the center of both ATAC and DHS regions (Extended Data Fig.3b,c). These findings strongly indicate the general preference of BRD4L for binding within nucleosome-free regions.

In contrast, an anti-FLAG ChIP-seq with N-terminal FLAG-tagged BRD4S showed a pair of peaks flanking the center of ATAC and DHS regions in MDA-MB-231 cells (Extended Data Fig.3b,c). This pattern is similar to the binding pattern of BRD4 revealed by the N-terminal antibody in MCF7 cells (Fig. 3h and Extended Data Fig. 3a), suggesting that BRD4S primarily binds at nucleosome-protected regions, as would be expected if bromodomain interactions played a major role in binding, and that the N-terminal BRD4 ChIP-seq mainly reflects the BRD4S binding. These results also show an unexpected role of the C-terminus of BRD4L, as it appears to play a critical role in the recruitment of BRD4L to nucleosome-free regions.

### BRD4 association with ER and Mediator is bromodomain-independent

ER and BRD4L (referred to as BRD4 hereafter) do not interact directly^25^, but co-immunoprecipitation (co-IP) of endogenous BRD4 revealed an estrogen-dependent ER-BRD4 association that remained unaffected by JQ1 (Fig. 4a). Co-IP of FLAG-tagged BRD4 fragments demonstrated that the bromodomains and a non-classic LXXLL sequence motif, which may enable ligand-dependent recruitment of co-activator by ER, are not required for BRD4 association with ER (Extended Data Fig.4a).

**Fig. 4.**
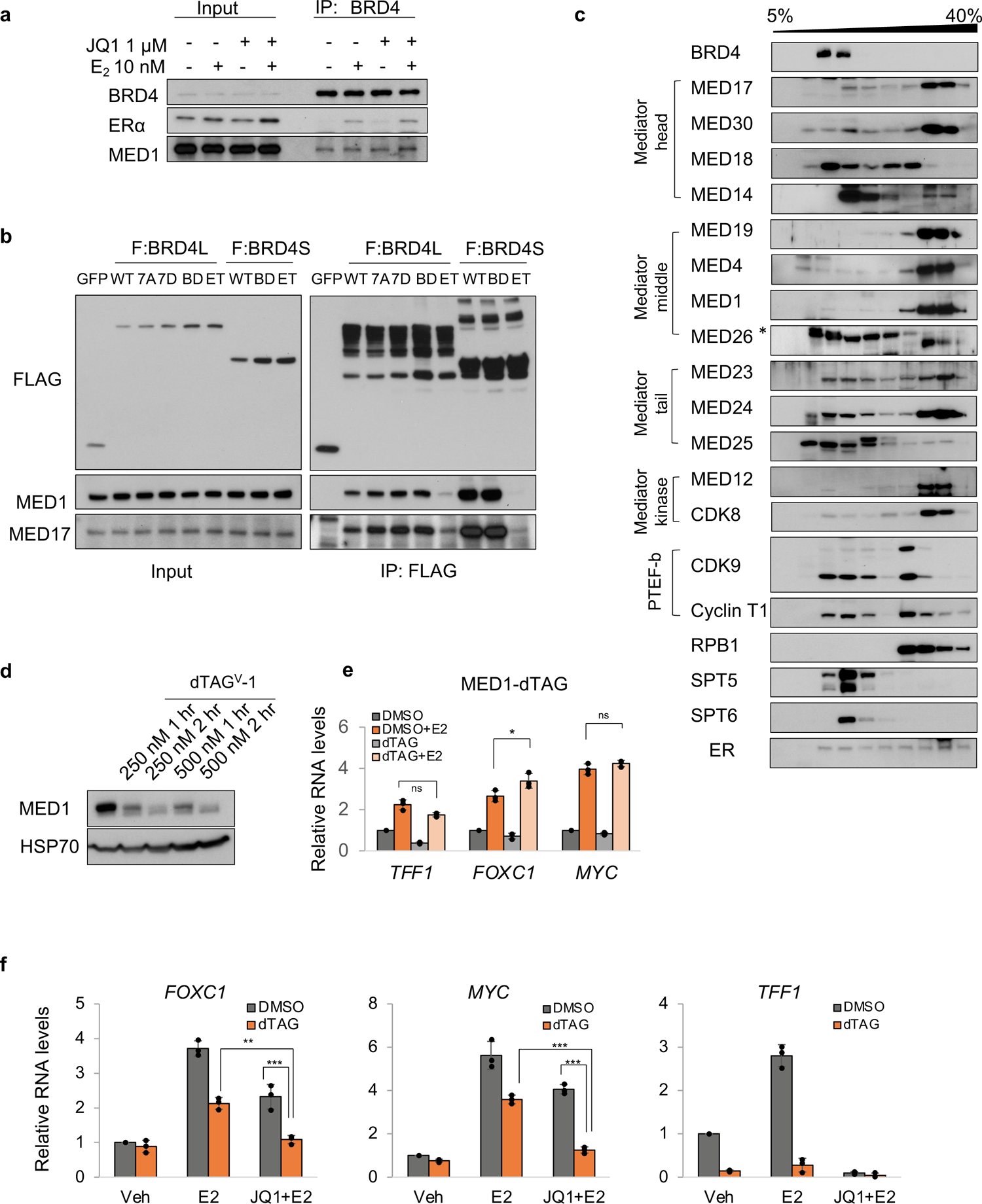
BRD4 associates with ER and Mediator. **a**, Immunoblots of indicated proteins from co-IP of BRD4 in MCF7 nuclear extracts. Cells were treated with or without JQ1 for 2 hours and then treated with or without E_2_ for 1 hour. JQ1 was also added to immunoprecipitates as indicated. **b**, Immmunoblots of BRD4L- and BRD4S-associated proteins in anti-FLAG immunoprecipitates of nuclear extracts from MCF7 cells transfected with the indicated plasmids. WT, wild type; 7A phosphorylation mutant; 7D phosphorylation mimic; BD, N140A/N433A mutant; ET, E651A/E653A mutant. **c**, Immunoblot of indicated proteins (left) from a 5-40% glycerol gradient of MCF7 nuclear extracts. Cells were grown in normal full medium. **d**, Immunoblot of MED1 protein levels in MED1-dTAG cells. A faint non-specific band exists below MED1. **e**, qPCR of gene expression changes in MED1-dTAG cells showing dispensable role of MED1 in ER-mediated transcription. **f**, qPCR of gene expression changes in MED14-dTAG cells upon dTAG^V^-1 treatment. Cells in **e** and **f** were treated with dTAG^V^-1 for 2 hours prior to E_2_ stimulation for 1 hour. Data shown as mean ± s.d. from triplicates. p values from two-tailed t-tests are depicted with asterisks. See also Extended Data Fig.4.

Next, we searched the literature for a factor that might have been reported to associate with BRD4 and act as an ER co-activator, which may explain the bromodomain-independent recruitment of BRD4. A recently proposed phase separation model suggests that MED1 plays a crucial role in ER-mediated transcription activation. MED1 possesses both LXXLL motifs that enable ligand-dependent recruitment of Mediator by ER and an intrinsically disordered region that compartmentalizes BRD4 and Pol II and enhances phase separation with ER^26–28^. Building on the reported MED1-BRD4 association in triple-negative breast cancer^29^, we hypothesized that BRD4 is recruited to ERBSs through an interaction with MED1/Mediator, which is recruited to chromatin-bound ER upon ligand activation, particularly at BETi-resistant genes. Co-IP of BRD4 from MCF7 nuclear extracts confirmed a constitutive association between BRD4 and MED1 that is independent of ER or bromodomains (Fig. 4a). To identify which domain of BRD4 is required for interaction with MED1, we expressed FLAG-tagged constructs of GFP, wild-type BRD4 (WT), mutants that eliminate (7A) or mimic (7D) CK2-mediated phosphorylation^29^, a bromodomain mutant (BD), and several truncations for co-IP experiments. WT BRD4 and all derived proteins except the 1-444 fragment, associated with MED1-containing Mediator as evidenced by probing IPs for MED1 and MED17, a core Mediator subunit (Extended Data Fig.4b). Therefore, either the middle region (442-722) or the disordered C-terminus of BRD4 is sufficient to associate with Mediator, while neither CK2-mediated phosphorylation nor the bromodomain is required for the association. The middle region contains the ET domain, which recognizes an amphipathic protein sequence motif^30,31^, such as a BRD9-like or WHSC1/L1-like short linear motif that is found in several Mediator subunits^7^. A BRD4 ET mutant (E651A/E653A), known to lose interaction with its binding proteins^30^, exhibited reduced association with MED1-containing Mediator in MCF7 cells (Fig. 4b). However, as observed with the BD mutant (above), the doxycycline-induced expression of the BD and ET double mutant fully restored *MYC* and *FOXC1* expression when the cells were treated with dBET6 (Extended Data Fig.4c). This likely reflects the residual Mediator association with the double mutant, which is sufficient for transcriptional activity for BETi-resistant genes. Given the widely presumed association of BRD4, MED1 and Pol II because of their colocalization on chromatin and in nuclear speckles with phase separation properties, we asked whether these factors associate within the same biological complex. In a glycerol gradient using MCF7 nuclear extracts (Fig. 4c), ER was evenly distributed in most fractions. In contrast, different Mediator subunits displayed somewhat distinct distribution profiles, with most of them sharing similar partition patterns. Although it has been reported that BRD4 interacts with the Mediator^3^, we found that BRD4 did not partition with the major fraction of Mediator or Pol II. This suggests that BRD4 is not a stable component of the Pol II-Mediator holoenzyme and that distinct Mediator (sub)complexes may exist for Pol II recruitment and BRD4 interaction.

### Mediator Disruption Impairs BRD4 Binding at ERBSs

Due to the complicated association between BRD4 and MED1/Mediator, it is challenging to disrupt this interaction through BRD4 mutations (as described above). Therefore, we asked if depleting MED1 would hinder the recruitment of BRD4 to ERBSs. Surprisingly, MED1 knockdown only slightly decreased *TFF1* transcription and failed to inhibit ER-dependent transcription of *FOXC1* and *MYC* (Extended Data Fig.4d,e). Similar results were observed when we completely depleted MED1 using dTAG-induced protein degradation, indicating that MED1 is not generally required for ER-mediated transcription in MCF7 cells (Fig. 4d,e and Extended Data Fig. 4f,g).

The dispensable role of MED1 suggested that both ER and BRD4 may interact with unidentified Mediator subunits. It is more likely that BRD4 interacts more generally with the Mediator complex, rather than solely relying on the MED1 subunit. To test this hypothesis, we performed dTAG-induced degradation of MED14, an essential architectural and functional component of the Mediator complex^32^. Despite successful engineering of MED14 in other cells, we failed to obtain a homozygous clone to completely degrade MED14, as it reportedly escapes X chromosome inactivation^33^. Nonetheless, in a heterozygous clone (Extended Data Fig.4h,i), dTAG treatment resulted in a partial loss of ER-dependent transcription of *MYC* and *FOXC1* and a complete loss of *TFF1* transcription (Fig. 4f). The combination of dTAG and JQ1 treatment showed a synergistic inhibitory effect on *MYC* and *FOXC1* transcription compared to dTAG or JQ1 alone. Subsequently, we attempted to disrupt Mediator function by depleting MED17, another critical subunit of Mediator^32^. Depleting MED17 using dTAG completely abolished transcription of *MYC* and *FOXC1,* irrespective of JQ1 treatment, further confirming a Mediator requirement for ER-dependent transcription (Fig. 5a-d). MED17 depletion also reduced the association of BRD4 with ER (Extended Data Fig.5a), consistent with intact Mediator serving as a bridge between ER and BRD4.

**Fig. 5.**
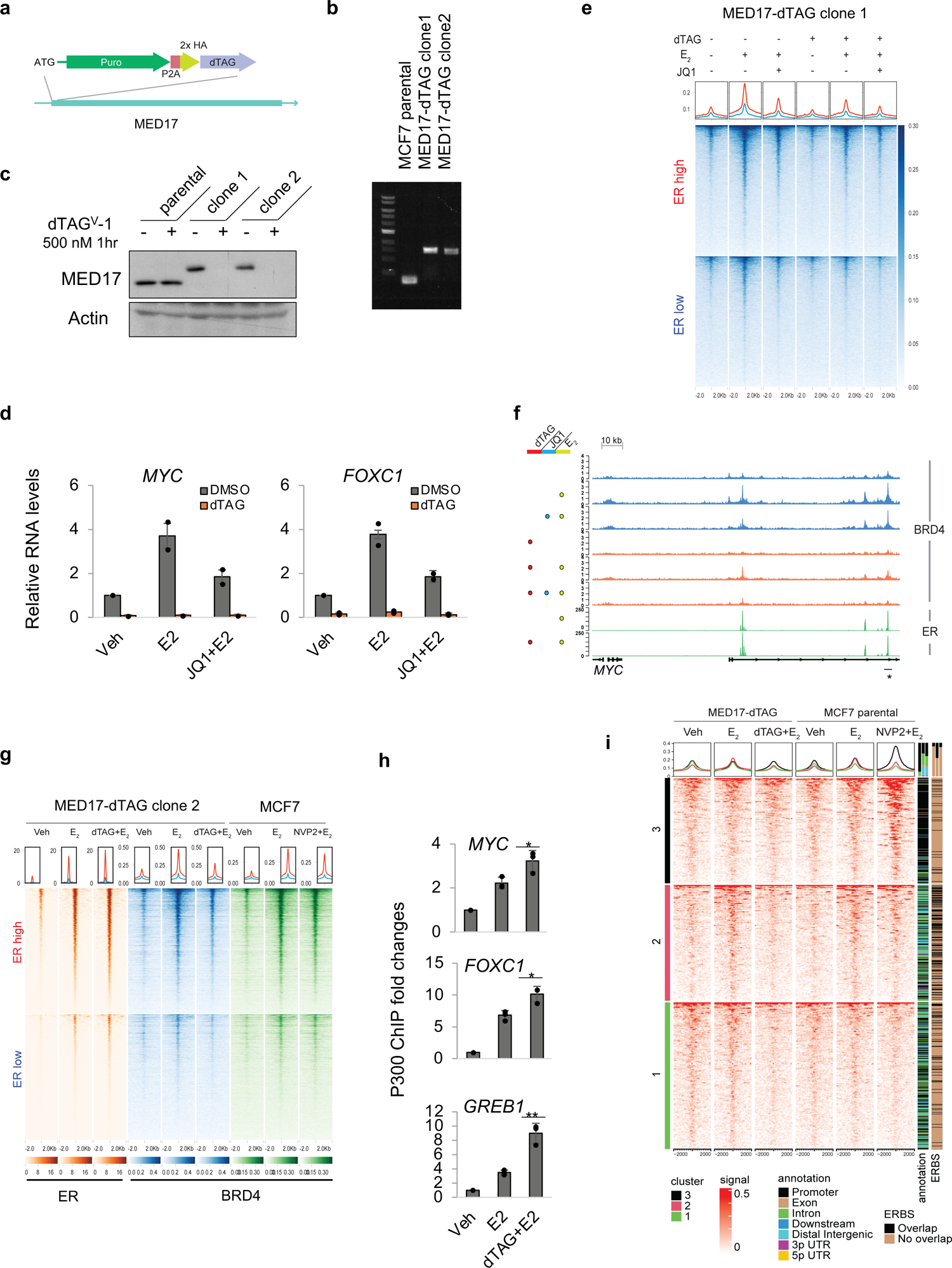
Mediator is required for BRD4 binding at ERBSs. **a**, Schematic of the knockin strategy for MED17-dTAG. **b**, Agarose gel electrophoretic analysis of PCR products from MED17-dTAG knockin cells. **c**, Immunoblot of MED17 in MCF7 (parental) or MED17-dTAG (clones 1 and 2) cells after indicated dTAG treatment. **d**, qPCR of *MYC* and *FOXC1* RNA levels in MED17-dTAG cells treated with or without 500 nM JQ1 or dTAG^V^-1 for 2 hours before E_2_ induction. Data shown as mean ± s.d. from two clones. **e**, Heatmap of BRD4 ChIP-seq signals at ERBSs from MED17-dTAG clone 1 with disuccinimidyl glutarate (DSG) crosslinking. Cells were treated with or without 100 nM JQ1 or 500 nM dTAG^V^-1 for 2 hours prior to E_2_ induction. **f**, ER and BRD4 ChIP-seq signals at the distal *MYC* enhancer (*) in MED17-dTAG cells under indicated conditions. **g**, Heatmap of ER and BRD4 ChIP-seq signals at ERBSs from MCF7 parental and derived MED17-dTAG clone 2 cells. Respective cells were treated with or without 250 nM NVP2 or dTAG for 2 hours prior to E_2_ induction. **h**, P300 ChIP-qPCR signals at gene enhancers in MED17-dTAG cells under indicated conditions. Data shown as mean ± s.d. from independent triplicates. p values from two-tailed t-tests are depicted with asterisks. **i**, Heatmap of BRD4 ChIP-seq signals at all BRD4 binding sites in MCF7 parental or MED17-dTAG cells (two clones combined). Respective cells were treated with or without dTAG or 250 nM NVP2 for 2 hours prior to E_2_ induction. See also Extended Data Fig.5.

Next, we analyzed BRD4 ChIP-seq signals at ERBSs upon MED17 depletion. Notably, MED17 depletion alone led to markedly reduced BRD4 binding at ERBSs, both with (Fig. 5e) and without disuccinimidyl glutarate (DSG) crosslinking (Extended Data Fig.5b). The combination of MED17 depletion and JQ1 treatment further decreased BRD4 binding, as exemplified both by heatmaps and by ChIP-seq signals at *MYC* and *GREB1* enhancers (Fig. 5e,f and Extended Data Fig.5c). We confirmed the diminished BRD4 binding by MED17 depletion at ERBSs in another clone (Fig. 5g and Extended Data Fig. 5d). To determine whether the reduction in BRD4 binding is a consequence of the loss of Mediator itself rather than ER binding, we performed ER ChIP-seq. In contrast to the reduced BRD4 binding, MED17 depletion increased ER binding at ERBSs, suggesting impaired ER turnover (Fig. 5f,g and Extended Data Fig. 5c). Coincident with increased ER binding, P300 binding was also enhanced at ERBSs upon MED17 depletion as measured by ChIP-qPCR (Fig. 5h). These results indicate that the decreased BRD4 binding was not due to reduced ER binding or P300-mediated acetylation and that Mediator plays a distinct role in BRD4 binding at ERBSs that is separate from the bromodomain-dependent interaction.

Since Mediator is required for transcription, we investigated whether the reduced binding of BRD4 could be a consequence of transcriptional inhibition. We performed BRD4 ChIP-seq following transcriptional inhibition by NVP2, a selective CDK9 inhibitor. At ERBSs, CDK9 inhibition resulted in a slight reduction of BRD4 binding (Fig. 5g and Extended Data Fig. 5d).

Interestingly, BRD4 binding was unaffected by NVP2 at *MYC* enhancers (Extended Data Fig.5d). Thus, although NVP2 led to a more severe transcriptional inhibition than did MED17 depletion, the change in BRD4 binding was less pronounced -- suggesting that attenuated BRD4 binding at ERBSs is not solely caused by transcriptional inhibition. The observed slight reduction in BRD4 binding might actually be a secondary effect of impaired transcriptional initiation and Mediator recruitment due to pausing^34^. Nonetheless, further comparison of the effects of MED17 depletion and CDK9 inhibition on BRD4 binding revealed noticeable differences. Unsupervised clustering divided all BRD4 binding sites into three major groups (Fig. 5i). Cluster 1 consisted of regions where BRD4 binding was unaffected by estrogen but decreased by MED17 depletion and modestly decreased by CDK9 inhibition. Cluster 2 included regions where BRD4 binding was enhanced by estrogen but decreased by MED17 depletion and modestly decreased by CDK9 inhibition. Cluster 3 comprised regions where BRD4 binding was unaffected by estrogen or MED17 depletion but increased by CDK9 inhibition. Cluster 2 was enriched with ERBSs, and cluster 1 likely contained enhancers without ERBSs, as clusters 1 and 2 showed a similar distribution across promoter, intron, and intergenic regions. However, cluster 3 primarily showed promoter occupancy. These results suggest that Mediator is necessary for optimal recruitment or stabilization of BRD4 at enhancers, whereas pausing primarily leads to the accumulation of BRD4 at promoters.

### BET depletion alters Pol II-associated protein profiles

A widely accepted model is that BRD4 activates elongation by recruiting CDK9, which in turn releases Pol II paused at promoter-proximal regions^9^. However, recent studies showed that depletion of BET proteins led to a global elongation defect without impaired CDK9 recruitment at active transcriptional start sites or active enhancers^10^. The elongation effect may be partially attributed to the intrinsic histone chaperone function of BET proteins, which requires bromodomains^5,6^. However, for BETi-resistant genes, there must be additional “effectors” at play that act independently of bromodomains to mediate the general elongation effect of BET proteins. To identify these putative effectors, we first re-assessed the controversial CDK9 recruitment by BET proteins. We examined the effect of BET depletion on CDK9 binding using ChIP-qPCR and included CDK9 inhibition by NVP2 as a positive control for pausing. At the *FOXC1* promoter, CDK9 occupancy was unaffected by pan-BET degrader dBET6 and enhanced by NVP2 (Fig. 6a and Extended Data Fig. 6a). At the *FOXC1* enhancer, both NVP2 and dBET6 actually enhanced CDK9 occupancy (Fig. 6b). Similar results were also observed at the major *TFF1* and *MYC* enhancers and the *MYC* promoter (Extended Data Fig.6b-e), suggesting that BET proteins, including BRD4, are not required for CDK9 recruitment. The *TFF1* promoter was not assessed because of the existence of a promoter-proximal enhancer. Surprisingly, we also observed an increase in ER binding by NVP2 and dBET6 treatments (Fig. 6b and Extended Data Fig. 6c), indicating that BET proteins are not necessary for the ER accessibility of ERBSs and that the increase in CDK9 enhancer binding may be a consequence of increased ER binding. Next, we examined Pol II levels at *FOXC1* and *MYC* promoters. At these promoters, both NVP2 and dBET6 further enhanced ER-dependent Pol II occupancy, suggesting that BET depletion led to promoter-proximal pausing (Fig. 6c and Extended Data Fig. 6e). As expected, both dBET6 and NVP2 reduced Pol II occupancy in the gene body (Extended Data Fig.6f). Moreover, NVP2 led to an accumulation of H2A.Z deposition, an H2A variant enriched at +1 nucleosomes, compared to the decrease in H2A.Z triggered by ER binding alone, likely reflecting E_2_-induced transcription that removes the +1 nucleosomes; and dBET6 partially reversed the ER-induced loss of H2A.Z, but to a lesser extent than NVP2 (Fig. 6d). These results indicate that, despite the fact that BET proteins are not required for P-TEFb recruitment, BET depletion partially resembles CDK9 inhibition -- it induces a pause state in the transcriptionally engaged promoter-proximal Pol II, leading to increased chromatin accessibility or reduced ER turnover at enhancers, retention of H2A.Z at +1 nucleosomes, and impaired Pol II processivity in the gene body.

**Fig. 6.**
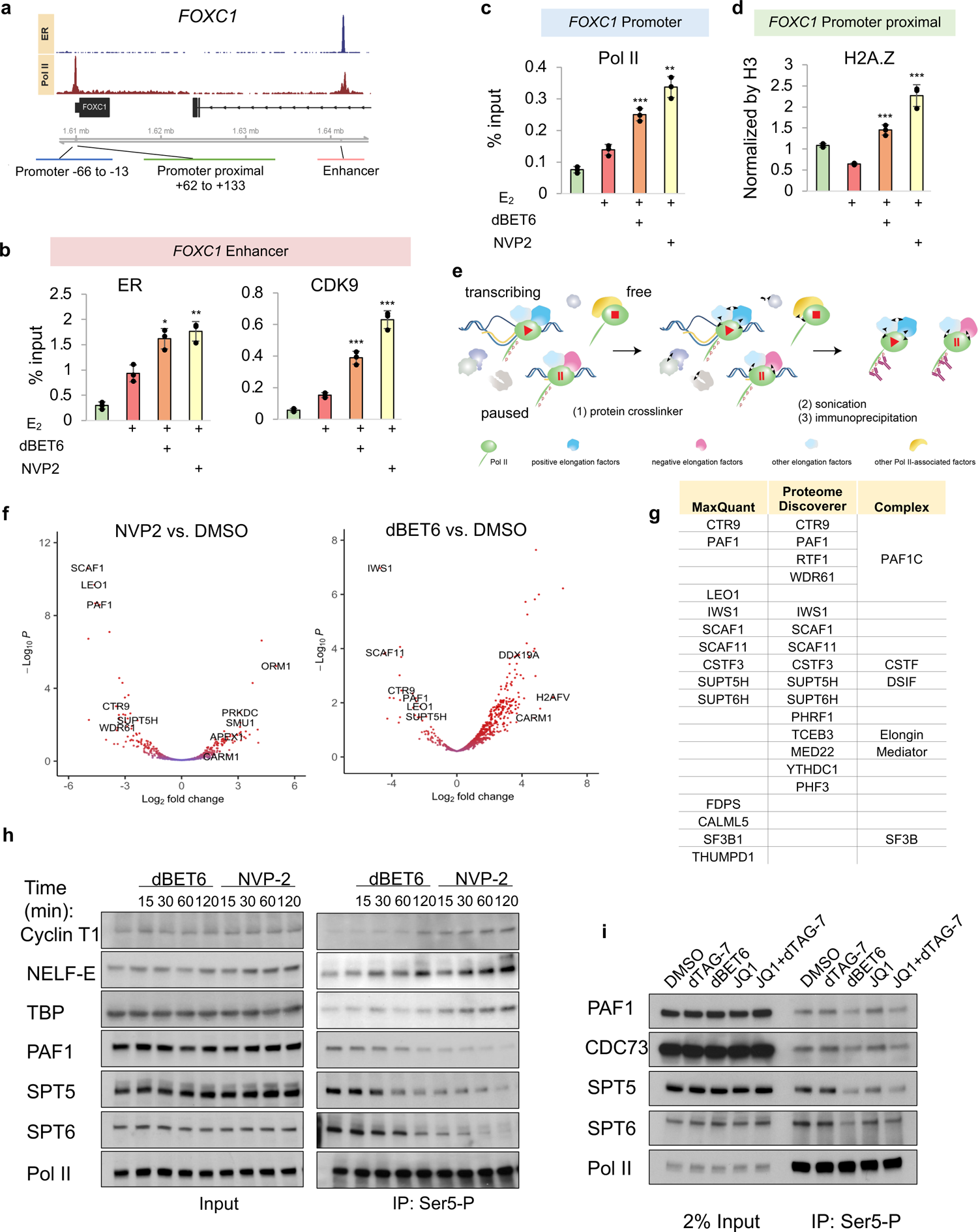
BET proteins regulate elongation factors association with Pol II. **a-d,** ChIP-qPCR analyses of indicated factors at different regions of *FOXC1* as illustrated in **a**. MCF7 cells were treated with or without NVP2 or dBET6 for 2 hours before E_2_ treatment. Data shown as mean ± s.d. from independent triplicates. p values from two-tailed t-tests (compared to E_2_ treatment alone) are depicted with asterisks. **e**, Diagram of workflow used to enrich post-initiation Pol II for label-free MS identification of proteins. Cells were lightly crosslinked by DTME and then subjected to sonication. The post-initiation Pol II complex was enriched using the Pol II CTD Ser5-P antibody. **f**, Volcano plot of differential protein interactions with post-initiation Pol II upon treatment of cells with 500 nM dBET6 or 250 nM NVP2 for 2 hours. Proteins identified by label-free MS. **g**, List of proteins showing significant loss of Pol II interactions in both NVP2 and dBET6 treatments by two (indicated) quantification methods. **h**, Immunoblot of indicated proteins from immunoprecipitation of Pol II CTD Ser5-P in MCF7 cells upon treatment with 250 nM NVP2 or 500 nM dBET6 for the indicated times. **i**, Immunoblot of indicated proteins in Pol II CTD Ser5-P immunoprecipitates from BRD4-dTAG cells upon treatment with NVP2, dBET6, JQ1 (500 nM) or dTAG7 for 2 hours. Cells in (**h**) and (**i**) were crosslinked and sonicated under the same conditions used in (**f**). See also Extended Data Fig.6.

CDK9 orchestrates the interactions of Pol II, negative elongation factor (NELF), and positive elongation factors (including DSIF, PAF1 complex, and SPT6) to enable transcription elongation^35^. However, little is known about the potential direct effect of BET proteins on the elongation complex. To address this, we utilized label-free mass spectrometry to specifically assess changes in the post-initiation Pol II complex. Cells were lightly crosslinked by dithiobismaleimidoethane (DTME), a reversible cysteine crosslinker, and the post-initiation Pol II complex was released from chromatin by sonication (Fig. 6e). Subsequently, the complex was enriched using a monoclonal antibody that recognizes Pol II CTD Ser5-P, which remains unchanged during transcription on chromatin^36^ and is largely unaffected by dBET6 or CDK9 inhibition^37^ (Fig. 6e, Extended Data Fig. 6g). This **<u>M</u>**ass **<u>S</u>**pectrometry with **<u>D</u>**TME **<u>C</u>**rosslinking (MSDC) strategy leverages Pol II Ser5-P, a natural marker of chromatin-associated Pol II, and significantly reduces background noise caused by formaldehyde crosslinking. We observed that DTME, unlike the lysine crosslinker DSP, does not introduce bias toward paused Pol II by CDK9 inhibition during Ser5-P enrichment (Extended Data Fig.6g). We compared the changes in the post-initiation Pol II complex associated proteins after 2 hours of dBET6 treatment and included NVP2 as a positive control for pausing, as well as IgG as a negative control. A total of 861 proteins were identified by MaxQuant in at least one condition across two independent replicates. Pol II Ser5-P enriched a similar number of proteins across different conditions, while IgG showed significantly lower protein enrichment (Extended Data Fig.6h). dBET6 treatment resulted in a decrease of 21 proteins and an increase of 122 proteins, with fold-changes exceeding 2-fold compared to DMSO, whereas NVP2 led to a decrease of 32 proteins and an increase of 13 proteins (p.adj<0.05, Fig. 6f). Among the 23 common proteins that exhibited significant changes in both conditions, many decreased proteins were transcription elongation factors that included SPT5, SPT6, and the PAF complex (PAF1C), which consists of CTR9, CDC73 (not consistently detected by MS), LEO1, PAF1, SKI8 (WDR61), and the dissociable subunit RTF1 (Fig. 6g). We also used the RPB1 subunit of Pol II to normalize protein abundance calculated by Proteome Discoverer and observed similar results, demonstrating a positive correlation between the changes induced by dBET6 and NVP2 relative to the control and a loss of elongation factors (Fig. 6g and Extended Data Fig. 6i). These findings suggest that the loss of BET proteins leads to a Pol II-associated protein profile that partially resembles Pol II pausing induced by CDK9 inhibition.

### BET proteins associate with elongation factors to regulate transcription

Given the results in the preceding section, we next focused on the elongation factors and validated the proteomic results from the crosslinking/immunoprecipitation approach (Fig. 6e) by Western blot. A remarkable decrease of Pol II-associated SPT5, SPT6 and PAF1 took place as early as 15 minutes after NVP2 treatment (Fig. 6h). By comparison, in the case of dBET6 treatment, a significant decrease of these protein associations occurred only after at least 60 minutes (Fig. 6h), probably because it took more time for a complete depletion of BET proteins. NVP2 and dBET6 treatments also reduced SPT5, SPT6 and components of PAF1C in the total cellular chromatin fraction (Extended Data Fig.6j), which supports the conclusion that the loss of Pol II-associated elongation factors was indeed a consequence of the lack of protein interactions and not differences in crosslinking efficiency. Although mass spectrometry failed to consistently detect all P-TEFb and NELF components, we observed an increased association of Cyclin T1 and NELF-E with Pol II upon NVP2 and dBET6 treatments (Fig. 6h and Extended Data Fig. 6k). These results are consistent with the enhanced association of CDK9 with chromatin as analyzed by ChIP-qPCR (Fig. 6b, Extended Data Fig. 6c,e) -- and further indicate that these treatments led to NELF-mediated pausing. Unlike NVP2 and dBET6, JQ1 alone had a much lower effect on SPT5 and SPT6 associations and barely any effect on PAF1C (Extended Data Fig.6k). A slight loss of PAF1C from the chromatin fraction was only detected when the JQ1 concentration was doubled (Extended Data Fig.6j). Despite a lack of apparent effects on PAF1C by individual JQ1 treatment or BRD4 depletion, the combination did reduce Pol II-associated PAF1C (Fig. 6i), suggesting that the chromatin association of PAF1C is less sensitive to JQ1 than SPT5 and SPT6. These results highlight a possible role for BET proteins in the regulation of elongation factor interactions.

An association between BET proteins and PAF1C has been reported^38,39^. Additionally, based on insensitivity to JQ1, our co-IP analyses revealed that all BET proteins associate with SPT5, SPT6, and PAF1C independently of known bromodomains interactions with acetylated histones (Fig. 7a), which may explain bromodomain-independent elongation regulatory functions. To determine whether the indicated elongation factors mediate the elongation effect of BET proteins, we individually depleted SPT5, SPT6 and PAF1 by shRNA-mediated knockdown or by dTAG-mediated degron. Knockdown of SPT5, which can either facilitate or (upon phosphorylation) release pausing, mildly enhanced ER-mediated transcription of *FOXC1* (∼ 4 kb) and *RET* (∼ 53 kb) (Extended Data Fig.7a,b), likely as a result of reduced pausing^36^. Knockdown of SPT6, a positive elongation factor, led to a slight decrease of ER-mediated transcription of *FOXC1* but not *RET* (Extended Data Fig.7c,d). PAF1 depletion by the dTAG approach led to a complete loss of PAF1, as well as destabilization of LEO1 and CTR9, within 2 hours (Fig. 7b).

**Fig. 7.**
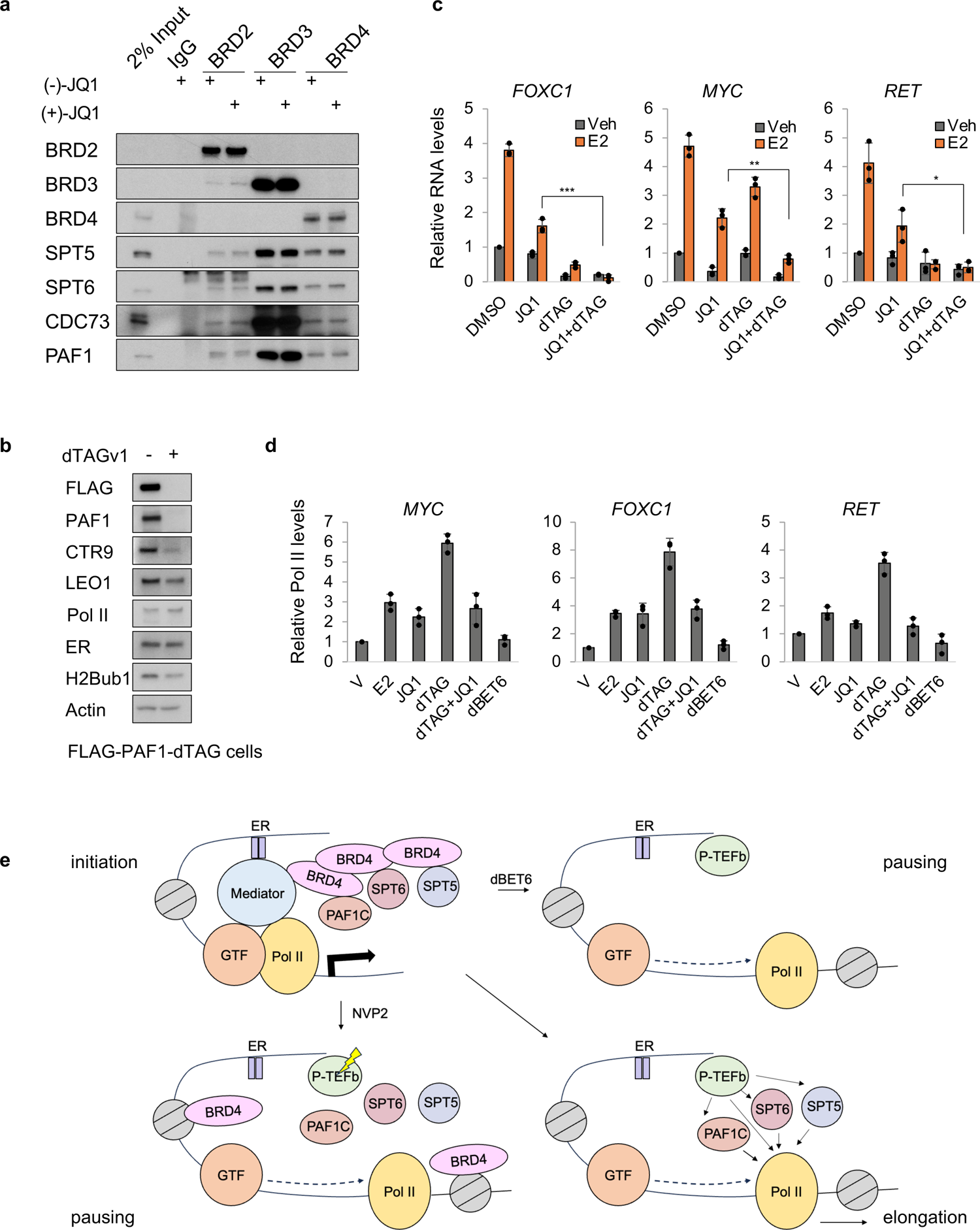
BETi-resistant gene transcription requires PAF1C. **a**, Immunoblot of indicated proteins from co-IP of BET proteins in MCF7 nuclear extracts. 5 µM of (-)-JQ1 or (+)-JQ1 was added during immunoprecipitation. **b**, Immunoblot of indicated proteins in FLAG-PAF1-dTAG cells treated with or without 500 nM dTAG^V^-1 for 2 hours. **c**, qPCR analyses of gene expression changes in FLAG-PAF1-dTAG cells treated with or without JQ1, dTAG or both prior to E_2_. **d**, ChIP-qPCR analyses of Pol II (NTD) binding at gene bodies in FLAG-PAF1-dTAG cells under indicated conditions. E_2_ was used in all treatments except for the vehicle group (V). Data shown as mean ± s.d. from independent triplicates. p values from two-tailed t-tests (compared to E_2_ treatment alone) are depicted with asterisks. **e**, Proposed model of BET proteins in ER-mediated transcription. Note that BRD2 and BRD3 are not shown for conciseness. They can also associate with elongation factors to facilitate transcription. GTF, general transcription factors. See also Extended Data Fig.7.

This loss of PAF1 impaired ER-mediated transcription of most tested genes (except *CA12*) and also sensitized their response to JQ1 inhibition (Fig. 7c Extended Data Fig. 7e), suggesting that PAF1 is required for most JQ1-resistant gene transcription. Notably, treatment with dBET6 significantly reduced the association of PAF1 with Pol II in the chromatin fraction (Extended Data Fig.7f). Consistent with previous observations of a slower transcribing Pol II in the gene body upon PAF1 deletion^40^, PAF1 depletion alone increased Pol II occupancy in the gene body (Fig. 7d). In contrast, JQ1 alone or in combination with PAF1 depletion did not significantly alter Pol II occupancy, while dBET6 reduced Pol II occupancy. These results suggest that Pol II occupancy is a mixed effect of different elongation factors by BET depletion, with loss of PAF1C displaying the most important functional consequences.

## Discussion

Adaptive resistance to BETi can emerge during BETi treatment through altered modifications of BET proteins or signaling pathways^29,41^. However, *de novo* failure of BETi to repress transcription remains unexplained. Here, we show that ER-mediated transcription of *MYC* and other oncogenes is intrinsically resistant to BETi due to (1) bromodomain-independent recruitment of BET proteins to enhancers and (2) bromodomain-independent recruitment of elongation factors by BET proteins to facilitate transcription (Fig. 7e).

### Bromodomain-independent recruitment of BRD4 requires Mediator

BETi-resistant chromatin association of BET proteins can occur through direct interaction with DNA-binding transcription factors, but ER is not the case. For such a bromodomain-independent recruitment, we have identified a multivalent interaction between BRD4 and Mediator that primarily accounts for the BRD4 occupancy at ERBSs. Using the dTAG approach, we have shown that bromodomains and Mediator recruit BRD4 to ERBSs independently and synergistically. The presence of BRD4 in a minor Mediator population, despite their frequent colocalization on chromatin, implies that the BRD4 and Mediator interaction may undergo dynamic regulation. Notably, hyper-phosphorylation of BRD4 was reported to promote a strong association of MED1^29^, but in our experimental conditions, neither BRD4 phosphorylation nor MED1 seems to affect the interaction between BRD4 and Mediator. Our results thus argue against the notion that MED1, with its strong phase separation properties, is generally required for ER-mediated transcription, although we cannot rule out the possibility that MED1 participates in the transcription of specific ER target genes.

Interestingly, it has been shown that bromodomain inhibition releases both Brd4 and Mediator subunits from enhancers of specific genes, including *Myc*, in a mouse leukemia cell line that depends on Med23, Med12 and Brd4^42^. These results indicate that bromodomain-independent recruitment of BRD4 by Mediator is specific to certain transcription factors that either use other factors to enhance/stabilize BRD4 binding or that selectively interact with the BRD4-containing Mediator. In contrast to results of our dTAG-mediated MED17 depletion, knockdown of Med23 and Med12 resulted in only a modest change or even an enhanced BRD4 occupancy^42^, as exemplified by results for the *Myc* superenhancer (Extended Data Fig.5e,f). Thus, the regulation of BRD4 and Mediator interactions/functions at enhancers is complex and likely context-dependent, and warrants further investigation.

In our experiments involving ectopic expression of BET mutants, bromodomain-mutated BRD4 fully rescued *MYC* transcription in BET-deficient (dBET6-treated) cells, whereas endogenous BRD4 failed to completely overcome the inhibitory effect of JQ1 on *MYC*. One possible explanation of these results is that the BRD4 mutant, which does not interact with JQ1, is not equivalent to the BRD4 engaged with JQ1. Another possibility is that BET proteins compete for the same effectors. Thus, we observed that when all endogenous BET proteins were depleted, the BRD4 mutant more efficiently restored *MYC* transcription. In this regard, it should be pointed out that bromodomain-independent effects of other BET proteins on transcription must also exist because the combination of BRD4 depletion and JQ1 did not fully recapitulate the impact of dBET6 on H2Bub1 and Pol II (Extended Data Fig.2d).

### BET proteins associate with elongation factors but do not recruit CDK9

Notably, we found that BET proteins associate with a common set of elongation factors. Although a previous study reported reduced levels of elongation factors on chromatin following BRD4 depletion^43^, the contribution of these factors to BET protein-mediated elongation was unclear; and it is known that effects of elongation factors are context-dependent^44^. Importantly, by individually depleting elongation factors, we determined that PAF1C, in particular, is functionally necessary for JQ1-resistant and ER-dependent transcription. Considering the association of BET proteins with elongation factors, we propose that BET proteins compact the distribution of these proteins near active genes to facilitate elongation, as a mechanism distinct from the known bromodomain-dependent BET chaperon activity. This may explain, at least partially, the BET protein contribution to elongation that is independent of CDK9 recruitment. It may also be noted that, at least in MCF7 cells, our results do not support a proposed role for BRD4 in P-TEFb recruitment ^45^, although it is possible that the C-terminus of BRD4 enhances P-TEFb activity^46^.

The recent study by Zheng et al.^45^ reported that the BRD4 mutant containing only the last 170 residues could functionally replace the intact BRD4 in P-TEFb recruitment, pausing release and gene expression except for genes like *MYC*, although the genomic localizations of the truncated mutant and CDK9 were not identified. This raised an interesting question as to whether the BRD4 C-terminus can act on its own as a co-activator as reported by Zheng et al.^45^. However, a prior study reported that a mutant encompassing only the last 189 residues of BRD4 actually exerted a dominant-negative effect on genes regulated by BRD4, suggesting that BRD4 requires both the N- and C-termini to function as a co-activator^47^. It is still possible that the truncated mutant behaves differently in different cell types. In a further analysis, we found that the key model genes (*BRD2, PRSS22* and *HSPA8*) used by Zheng et al. showed no corresponding changes in gene expression from their RNA-seq results^45^, despite significant alterations in Pol II and Cyclin T1 binding profiles when BRD4 was depleted or rescued by bromodomain-less BRD4 mutants. These conflicting results could be attributed to the fact that the authors generated different cell lines expressing various BRD4 mutants under the control of Tet transactivator and then compared different cells only in the presence of doxycycline without considering different basal levels or activities of the investigated factors in different cell lines.

### MSDC is a simple approach to investigate Pol II post-initiation complexes

Beyond our elucidation of transcriptional regulatory mechanisms by BET proteins, we have developed a new strategy for proteomic analysis of Pol II-associated proteins that, due to its greater specificity, provides a more detailed understanding of changes in the post-initiation Pol II complex compared to the widely used chromatin-mass spectrometry (chromatin-MS) and ChIP-MS methods. For instance, upon dBET6 treatment, we observed a significant 80-91% loss of PAF1C components from Pol II by this method, whereas the chromatin-MS method only detected an approximate 30% loss of PAF1C components from chromatin^43^.

From a transcriptional perspective, our study reveals bromodomain-independent mechanisms underlying BET protein recruitment and function in ER-mediated transcription. Elucidation of the mechanism of ER transcriptional resistance to BETi provides a model to study similar events of other transcription factors. Combining BETi with other targeted approaches, such as BRD4 depletion or inhibition of other bromodomain-independent BRD4 interactions, may allow for use of lower concentrations of BETi to achieve therapeutic efficacy and thus to reduce adverse effects.

## Acknowledgments

We thank laboratory members Keiichi Ito, Sohail Malik and Takashi Onikubo for their comments on the manuscript, the Proteomics Resource Center (RRID:SCR_017797) and the Flow Cytometry Resource Center (RRID:SCR_017694) at The Rockefeller University. This work was supported by NIH grants CA234575 and CA273709 to R.G.R. and by the Robertson Therapeutic Development Fund and the Shapiro-Silverberg Fund for the Advancement of Translational Research to S.Z. S.Z. was supported by an AACR-John and Elizabeth Leonard Family Foundation Basic Cancer Research Fellowship.

## Authors’ Contributions

S.Z. conceived the project, designed and performed experiments, analyzed and interpreted data, and wrote the manuscript. R.G.R. supervised the project and wrote the manuscript.

## Declaration of interests

Both authors declare no competing interests.

## Methods

### Cell lines and culture methods

MCF7 cells (a gift from Dr. Sarat Chandarlapaty, Memorial Sloan Kettering Cancer Center) and MCF7/FRT/TR cells (a gift from Dr. Reuven Agami, Netherlands Cancer Institute) were maintained in DMEM/F12 medium (Thermo Fisher 11330-032 or GenClone 25-503) with 5 % fetal bovine serum (R&D systems, S11150). HEK 293T cells from ATCC were maintained in DMEM medium with 10% FBS. All cells were grown at 37°C and 5% CO2 free of mycoplasma. To analyze ER-dependent gene activation or other effect of estrogen, the cells were cultured for 3-4 days in phenol red-free DMEM/F-12 (Thermo Fisher 11039021) supplemented with 5% charcoal-dextran-treated FBS. 17-β-estradiol (Sigma-Aldrich) was used at 10 nM in all experiments.

### Inhibitors and antibodies

dBET6 (#HY-112588), NVP-2 (#HY-12214A) were purchased from MedChemExpress. JQ1 (#A1910) was purchased from ApexBio. THZ1 (#9002215), (−)-JQ1 (#11232), MZ1 (#21622), ABBV-744 (#30470), I-BET762 (#10676), fulvestrant (#10011269) were purchased from Caymen. THZ531 (#AOB8107), GSK778 (#AOB11459), GSK620 (#AOB11616) were purchased from AOBIOUS. dTAG-7 (#6912), dTAG^V^-1 (#6914) were purchased from Tocris. Primary antibody for CDC73 (#66490-1-Ig) was purchased from Proteintech. FLAG antibody (#F1804) was from Sigma. ERα (#D8H8, WB), H2Bub1 (#5546), MED26 (#13641), and Rpb1 NTD (#D8L4Y, ChIP) antibodies were from Cell Signaling. PAF1 (#172A), LEO1 (#A300-174A), CTR9 (#A301-395A), RTF1 (#A300-179A), BRD4 (#A700-004, WB; IP), BRD4 (# A301-985A100, ChIP), BRD3 (#A700-069, IP), BRD2 (#A302-583A, WB, IP), MED1 (#A300-793A), MED24 (#A301-472A), and SPT6 (#A300-801A) antibodies were from Bethyl Laboratories. Pol II CTD Ser2P (#3E10) and Ser5P(#3E8) antibodies were from Activemotif. SKI8 (#OACA06119) and MED17 (#OAGA04959) antibodies were from Aviva Systems Biology. RPB1 (F12, N20, WB; IP), Cdk9 (D-7), cyclin T1 (E-3), NELF-E (F-9), BRD3 (2088C3a, WB), BRD2 (G4, WB), SPT5 (D-3), Erα (F-10, D-12, ChIP), U1 snRNP 70 (C-3), CDK8 (C19), P300 (N15), HSP70 (W27), MED18 (E14), NELF-A (A-20), and ß-Actin (C4) antibodies were from Santa Cruz. H2A.Z (ab4174), BRD4 (ab128874, NTD, WB), H3K27Ac (ab4729), H3K4me1 (ab8895), H3K4me3 (ab8580), MED4 (ab129170), Histone H3 (#ab1791), MED25, and MED23 (ab200351) antibodies were from Abcam. MED14 (A04799-2) antibody was from Boster. Antiserum for TBP, MED12, MED30, SPT16, RPB6 were made inhouse. Antiserum for MED19 was a kind gift from Conaway lab.

### Plasmids for gene expression

Constructs for BRD4 fragments p6345 MSCV-CMV-Flag-HA-Brd4 1-722, p6347 MSCV-CMV-Flag-HA-Brd4-444-722, p6348 MSCV-CMV-Flag-HA-Brd4 1047-1362, p6349 MSCV-CMV-Flag-HA-Brd4 1224-1362, p6346 MSCV-CMV-Flag-HA-Brd4-1-444 were from Peter Howley (Addgene 31352, 32886, 31353, 31354, 31355). Constructs for pcDNA5-Flag-BRD4 WT, BD, 7A, 7D were from Kornelia Polyak (Addgene plasmid # 90331, 90006, 90005, 90007). pcDNA5-Flag-BRD4-ET was altered by mutagenesis to harbor E651A/E653A point mutations. pcDNA5-Flag-BRD4S-WT, -BD, and -ET were made from the constructs with same mutations in long isoforms. For PAF1-dTAG, the CMV promoter was removed from pLVX-IRES-hyg vector with ClaI and XhoI digestion. The natural promoter and the 5’ UTR of *PAF1* and full cDNA sequences were cloned into the vector. The PAF1 cDNA was cloned from MCF7 cells using the following primers: forward-ATGGCGCCCACCATCCAGAC, reverse-GTCACTGTCACTATCAGCTTC. The promoter sequence was cloned from MCF7 genomic DNA using the following primers: forward - TAGCGCTCTCTCCGACTAAC, reverse-AGAGCTCCAGCGAGACTCA. The cloned sequences were verified by Sanger sequencing. Promoter, 5’ UTR, and 3xFLAG sequences were then synthesized for insertion before the start codon of PAF1, and the full plasmid without dTAG was assembled using In-Fusion® HD Cloning Kits (Takara). A second round of PCR was used to amplify the inserted sequence, which was assembled with dTAG by In-Fusion® HD Cloning Kits to create PAF1-dTAG with 3xFLAG at the N-terminus and dTAG at the C-terminus. Primer sequences are listed in Supplementary Table 1.

### Inducible expression of BET mutants

In experiments using the Flp-In system, BET mutants were cloned into pcDNA5/FRT/TO and co-transfected with pOG44, which encodes Flp recombinase, into MCF7/FRT/TR cells. Transfected cells were selected for hygromycin resistance for 2 weeks. Protein expression was induced by overnight treatment with 100 ng/ml doxycycline. pCDNA5-Flag-BRD4-BD mutant was from Kornelia Polyak (Addgene 90005). Wild-type BRD2 and BRD3 from GFP-BRD2 (Addgene 65376) and GFP-BRD3 (Addgene 65377) from Kyle Miller were cloned into the pcDNA5/FRT/TO vector. Then, BRD2-BD mutant (N156A/N429A) and BRD3-BD mutant (N116A/N391A) were made using QuikChange XL Site-Directed Mutagenesis Kit (Agilent Technologies 200517-4). The BRD4-BE mutant was modified from BRD4-BD to harbor E651A/E653A point mutations.

In Tet-inducible expression experiments using Tet transactivator, an IRES sequence was inserted between the TRE3GS promoter and EGFP from the XLone-GFP construct from Xiaojun Lian (Addgene 96930) to allow EGFP expression as a reporter. BET mutants were then cloned upstream of IRES under the control of the TRE3GS promoter. XLone-derived constructs were co-transfected with a plasmid encoding piggybac transposase into MCF7 cells with TransIT-LT1 Transfection Reagent (Mirus). Transfected cells were selected for blasticidin resistance for 2 weeks. Protein expression was induced by overnight treatment with doxycycline at 10 ng/ml.

### RNA analysis by quantitative real-time PCR (RT-qPCR)

TRIzol reagent (Thermo Fisher 15596026) was added to cells grown in six-well plates to isolate total RNA according to the manufacturer’s protocol. One or one-half microgram of total RNA was reverse-transcribed using iScript gDNA Clear cDNA Synthesis Kit (Bio-Rad 1725035). If RNA was treated with TURBO DNase (Thermo Fisher AM2238), RNA was then purified with TRIzol or Phenol:Chloroform:Isoamyl Alcohol 25:24:1 (Sigma P3803) and reverse-transcribed using cDNA Synthesis Kit without additional DNase treatment. Quantitative real-time PCR using Powerup SYBR Green PCR Master Mix (Thermo Fisher A25741) or QuantiTect SYBR Green mix (QIAGEN) was performed on a 7300 or QuantStudio 3 Real-time PCR System (Applied Biosystems). The annealing/extension temperature was 60°C for all primer sets listed in Supplementary Table 2. Relative expression levels were calculated using the delta-delta Ct method with GAPDH mRNA for short-term experiments or 5S ribosomal RNA for long-term experiments (BETi incubation overnight) as reference genes. 17-β-estradiol was added for 1 hour to measure ER-dependent transcription, if not stated otherwise.

### Immunoprecipitation

#### For endogenous proteins

For immunoprecipitation without crosslinking, nuclear pellets collected from one 10 cm dish were suspended in 0.3 ml buffer B (20 mM Tris pH 7.5, 25% glycerol, 0.42 M NaCl, 1.5 mM MgCl_2_, 1 mM KCl, 0.5% NP40, 0.2 mM EDTA, 0.5 mM DTT, sodium butyrate, protease and phosphatase inhibitors) and incubated for 15 min on ice. After centrifugation at 13,000 rpm for 15 min at 4°C, supernatant was collected and diluted with 0.6 ml buffer A (10 mM Tris pH 7.5, 1.5 mM MgCl_2_, 10 mM KCl, 0.5 mM DTT, sodium butyrate, protease and phosphatase inhibitors). The nuclear extracts were then incubated at 4 °C overnight with 1 μg antibody. The following day, immunoprecipitates were incubated with Dynabeads Protein G or A at 4 °C for 2 h. Beads were washed six times with buffer B containing 150 mM NaCl and 0.5% NP-40 and boiled in loading buffer. For immunoprecipitation of crosslinked Pol II, cells were collected from 10 cm dishes and sonicated in X-IP buffer (10 mM Tris-HCl pH 7.5, 150 mM NaCl, 1 mM EDTA, 1 mM EGTA, 1% NP-40, 0.5% sodium deoxycholate). The lysates were cleared by centrifugation and incubated at 4 °C overnight with Dynabeads Protein G pre-coated with Pol II CTD Ser5-P antibody. Beads were washed six times with X-IP buffer and boiled in loading buffer. For mass spectrometry, beads were washed six times with IP-MS buffer (10 mM Tris-HCl pH 7.5, 1% NP-40, 150 mM NaCl) and six times with PBS.

#### For exogenous proteins

For each 10 cm dish of cells, 5 µg of plasmid was used for transfection and the cells were allowed to grow for two days before collection for nuclear extract preparation.

### Cross-linking and mass spectrometry

Cells growing exponentially in 10% FBS were treated with DMSO, 500 nM dBET6 or 250 nM NVP2 for 2 hours and then washed twice with PBS at room temperature. DTME (dithiobismaleimidoethane, Thermo Scientific) or DSP (dithiobis(succinimidyl propionate), Thermo Scientific) was dissolved in DMSO at 50 mM and quickly added to PBS at 0.5 mM final concentration to make the cross-linker solution. Cells were incubated with the cross-linker solution for 30 minutes at room temperature, washed twice with PBS and collected using cell lifters. Samples were resuspended, sonicated and centrifuged for immunoprecipitation. Immunoprecipitates from three tubes of the same conditions were combined for on-bead trypsin digestion. Label-free quantitative mass spectrometry was performed by the Proteomics Resource Center at The Rockefeller University. Two independent replicates were processed independently. Proteins identified by ProteomeDiscoverer v. 1.4.0.288 (Thermo Scientific) were filtered by IgG condition to remove background, which resulted in 1377 remaining proteins. These proteins were further filtered by PSM>2 and identification in both replicates under at least one condition. Missing values were imputed by the number of 100000. The remaining 668 proteins were normalized by RPB1 abundance. Fold changes (log2) were calculated by transformed protein abundance under treated conditions relative to DMSO. Fold changes (log2) were considered biologically significant if the changes of replicates were unidirectional and the absolute mean was more than 1. Protein quantitation was also generated by MaxQuant. iBAQ values were used as input for DEP v1.14.0^48^. Proteins were filtered for those that were identified in both replicates of at least one condition. Missing values were imputed by “MinDet” and q=0.01. For the remaining 861 proteins, changes were considered significant if the log2 fold changes relative to DMSO were more than 1 and padj< 0.05.

### Chromatin IP and ChIP-sequencing

Cells were grown in 15 cm dishes for 3-4 days in phenol red-free DMEM/F-12 supplemented with 5% charcoal-dextran-treated FBS. Cells were treated with DMSO, 500 nM JQ1 and/or 500 nM dTAG for 2 h prior to incubation with 10 nM estrogen. After estrogen treatment for 40 min, cells were fixed with 1% formaldehyde at room temperature for 10 min. Glycine was then added to quench cross-linking. Fixed cells were washed twice with PBS, collected and resuspended in cold lysis buffer 1 (50 mM HEPES–KOH, pH 7.5, 140 mM NaCl, 1 mM EDTA, 10 % glycerol, 0.5 % Igepal CA-630, 0.25 % Triton X-100, 1× protease inhibitor (Roche)). After 10 min rotation in a cold room, cells were centrifuged and resuspended in cold lysis buffer 2 (10 mM Tris–HCl, pH 8.0, 200 mM NaCl, 1 mM EDTA, 0.5 mM EGTA, 1× protease inhibitor). After 10 min rotation, the pellets were collected and sonicated by probe (Branson sonifier) in 300 μL lysis buffer 3 (10 mM Tris–HCl, pH 8.0, 100 mM NaCl, 1 mM EDTA, 0.5 mM EGTA, 0.1 % Na-deoxycholate, 0.5 % N-lauroylsarcosine, 1× protease inhibitor) to achieve fragment sizes below 700 bp. Insoluble materials were removed by addition of 30 μl 10 % Triton X-100 followed by centrifugation. Chromatin lysates were either incubated with antibody or antibody-coated magnetic beads overnight with rotation in the cold room. On the next day, beads were washed six times in RIPA buffer (50 mM HEPES–KOH, pH 7.5, 500 mM LiCl, 1 mM EDTA, 1 % NP-40 or Igepal CA-630, 0.7 % Na-deoxycholate) followed by one time in TE buffer. After that, beads were resuspended in 50 mM Tris–HCl, pH 8.0, 10 mM EDTA, 1 % SDS for reverse cross-linking at 65 °C overnight. RNA and proteins were removed by RNase A treatment at 37 °C for 30 min and Proteinase K digestion at 55 °C for 1 h, respectively. DNA was purified using QIAquick PCR Purification Kit (QIAGEN). For ChIP with H2A.Z and H3 histones, crosslinked cells were incubated with PBS containing 0.5% Triton X-100 for 3 min on ice. Nuclei were pelleted by centrifugation, washed and resuspended in MNase digestion buffer (10 mM Tris–HCl, pH7.4, 15 mM NaCl, 60 mM KCl, 0.15 mM spermine, 0.5 mM spermidine, 1 mM CaCl_2_). 0.03 μL MNase (New England Biolabs) was added per 100 μL mixture to digest chromatin at 37°C for 5 min and stopped by addition of lysis buffer 3. The mixture was briefly sonicated and processed as above. Eluted DNA was either used for quantitative real-time PCR or for library preparation. In some ChIP samples used to investigate the role of Mediator, chromatin from mouse pre-adipocytes was mixed in as spike-in control, but not used for calculation because of low enrichment in mouse cells.

Double crosslinking was used for FLAG-BRD4^mut^ and some MED17 depletion experiments (Fig. 3f and 5e). JQ1 was added to 100 nM instead of 500 nM to evaluate the combinatory effect of JQ1 and MED17 depletion. DSG was dissolved in PBS at 1 mM and added to PBS-washed cells for 30 min at room temperature. The remaining procedures were the same as above.

ChIP-seq libraries were prepared with NEBNext Ultra™ II DNA Library Prep with Sample Purification Beads (New England Biolabs) and sequenced by BGI. Pair-end reads were aligned to the human genome (hg38) with decoy sequences using Bowtie2 v2.4.4^49^. Samtools v1.13^50^ was used to generate BAM files for peak calling with MACS3 v3.0.0a6^51^. ChIP signals were normalized to counts per million reads by deepTools2.0 v3.5.1^52^ with duplicates ignored. Regions overlapping with ENCODE Blacklist regions were excluded from analysis. Peak annotation was performed with ChIPseeker^53^. Heatmaps were generated by deepTools or profileplyr^54^. Galaxy was also used in some experiments. For analysis using Galaxy, all procedures were performed identically to the above process except for some software versions that were preinstalled by Galaxy^55^.

### Analysis of public datasets

Gene expression profiles of MCF7 cells in response to estrogen and JQ1 were analyzed based on RNA-seq from GSE109570 and GSE55922. Gene expression was quantified by Salmon v1.5.2^56^ on the hg38 reference genome. Differential analysis was performed with DESeq2^57^ and visualized by heatmap with unsupervised clustering using pheatmap v1.0.12.

ER binding sites were derived from GSM365926 and GSE59530. ER ChIP-seq signals from GSM365926 was used to divide common ER binding sites into ER-high and ER-low sites. Raw reads of BRD4 ChIP-seq using an N-terminal antibody were processed from GSE55921 through Galaxy. Raw reads of BRD4 ChIP-seq in mouse leukemia cells were processed from GSE78221 through Galaxy. Reads from human and mouse samples were aligned to hg38 female and mm10, respectively. Processed ChIP-seq data of MED1 (GSE125594), H3K27ac (GSE45822), H4K12ac (GSE65886), Pol II (GSM365930) and DNase-seq data (GSM1255280) from MCF7 cells were downloaded from NCBI. Gro-seq data were from GSM1115995. ChIP-seq and DNase-seq data from MDA-MB-231 cells were from GSM2242136 and GSE136151. ATAC-seq from MCF7 and MDA-MB-231 cells were from GSM2572565 and GSE216496, respectively.

### Glycerol gradient centrifugation

Nuclear extracts from MCF7 cells growing in regular medium were dialyzed in 300 ml dialysis buffer (20 mM Tris pH 7.5, 0.15 M NaCl, 0.75 mM MgCl_2_, 0.5 mM KCl, 0.5% NP40, 0.2 mM EDTA, 0.5 mM DTT and PMSF) for 3 hours. 200 μl of extract was layered on top of a 5 to 40% (v/v) glycerol gradient in 3.8 ml dialysis buffer and centrifuged at 38,000 rpm (SW 60) for 16 hours. 200 μl fractions were collected slowly from the top of the gradient tube.

### Cell viability

Cells were seeded in medium with 5% FBS at 2000 cells per well in 96-well plates. The next day, medium was replaced with fresh medium containing 10% FBS and BET inhibitors or other indicated chemicals. Cells were allowed to grow six days and medium was changed on the third day. Relative cell numbers were measured by BMG LABTECH plate reader using Cell Counting Kit-8 (dojindo, CK04) according to the manufacturer’s protocol.

### Lentivirus Production and Transduction

#### For gene knockdown

shRNA sequences targeting MED1 or SPT6 were ligated to Tet-pLKO-puro to mediate MED1 or SPT6 knockdown. shRNAs for GFP, SPT5, BRD2, BRD3, and BRD4 were purchased from Sigma or Dharmacon or generated in pLKO.1 vector. Scramble shRNA was from David Sabatini (Addgene 1864). Sequence information is listed in Supplementary Table 3. The lentiviral vectors were co-transfected with packaging vectors psPAX2 and pMD2G (Addgene 12260 and 12259) into 293T cells for lentivirus production. MCF7 cells were transduced by unconcentrated lentiviruses with polybrene (6 μg/ml). The next day, cells were replenished with fresh medium. After 72 hr of transduction, cells were selected with 0.5 μg/ml puromycin for 6 days. For shMED1 Tet-on cells, doxycycline was added to complete medium at 100 ng/ml for 3 days before cells were transferred to phenol-red free medium with charcoal-dextran-treated FBS. For shSPT6 Tet-on cells, doxycycline was added when cells were transferred to phenol-red free medium.

#### For gene expression

For FLAG-PA1-dTAG expression, after lentiviral production and transduction, cells were selected with 100 μg/ml Hygromycin B for 6 days. For ESR1-Y537S expression, cells positively transduced by virus bearing pHAGE-ESR1-Y537S (Addgene 116374 from Gordon Mills & Kenneth Scott) were selected by GFP at The Flow Cytometry Resource Center at The Rockefeller University and maintained in full medium with fulvestrant (0.5 nM).

### Target-specific protein degradation

An endogenous knock-in of dTAG-MED17 was generated by CRISPR/Cas9-mediated PITCh (Precise Integration into Target Chromosome) system^58^. Plasmids for endogenous knock-in of dTAG-MED1 and dTAG-MED14 were kind gifts from Georg E. Winter (CeMM Research Center for Molecular Medicine of the Austrian Academy of Sciences, Austria). For dTAG-MED14, we failed to get homozygous knock-in clones from over 40 clones. We replaced the BSD cassette with Puro, but we still failed. Plasmids for dTAG-BRD4 were from James Bradner & Behnam Nabet (Addgene 91792 and 91794). MCF7 cells were plated in 10 cm plates and co-transfected with 2 µg of pCRIS-PITChv2-donor plasmid and 4 µg of pX330A-gRNA using TransIT-LT1 Transfection Reagent. After 2 days, transfected cells were replenished with fresh medium containing puromycin or blasticidin for selection. After a week, single cells were manually cultured in 96-well plates to expand to visible clones. Viable clones were genotyped by PCR and protein degradation efficiency was verified by immunoblotting. PAF1-dTAG cells were generated by knocking out endogenous *PAF1* using CRISPR/Cas9 and adding back FLAG-PAF1-dTAG using lentiviruses.

### Quantification and statistical analysiss

Data was analyzed with the statistical test described in the corresponding figure legend. Statistical significance was depicted as the p-value for single comparisons: ∗ <0.05, ∗∗ <0.01, ∗∗∗ <0.001, ns >0.05.

## Data availability

All ChIP-seq data have been deposited at GEO repository under the accession number GSE241695.

## Supplementary figure legends

**Extended Data Fig. 1.**
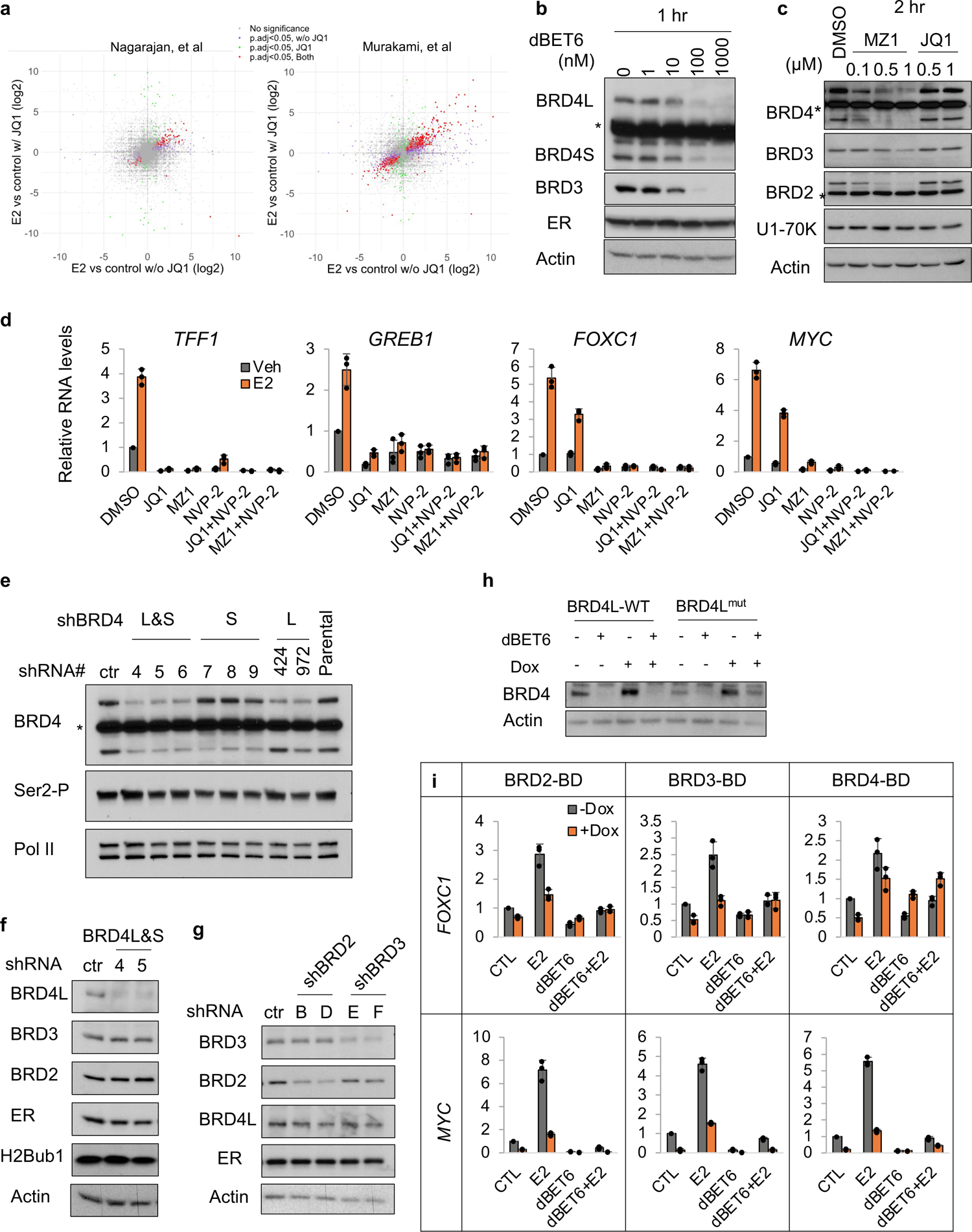
Some ER target genes are resistant to bromodomain inhibition but not BET depletion, related to **Fig. 1**. **a**, Effect of E_2_ on gene expression with or without JQ1 treatment in MCF7 cells. Data represent log2 fold change in comparison of E_2_ with vehicle treatment with or without JQ1 pre-treatment. Color shows statistical significance in the condition without JQ1 (purple), with JQ1 (green) or both settings (red). **b,c,** Immunoblot of indicated proteins with doses of dBET6, JQ1 or MZ1 treatments. The asterisk in **b** indicates an unknown band detected by the Abcam antibody that recognizes the N-terminus of BRD4. Cells in **b** and **c** were cultured in full medium (5% FBS, phenol-red). The asterisks in **c** indicate unknown bands. **d**, Effect of JQ1 (1 μM), MZ1 (500 nM) and NVP2 (250 nM) on ER-mediated gene transcription measured by qPCR. MCF7 cells were treated with inhibitors or degraders, as indicated, for 2 hours prior to a 2-hour induction by E_2_ (10 nM). NVP2 is a CDK9 inhibitor that serves as a positive control. **e-g**, Immunoblot of indicated proteins in whole cell lysates of MCF7 cells expressing shRNAs for GFP (ctr), BRD4-L (424, 972), BRD4-S (7, 8, 9), BRD4 (4, 5, 6), BRD2 (B, D), or BRD3 (E, F). The asterisk indicates an unknown band. **h**, Immunoblot of BRD4 expression in MCF7 FRT/Tet-on whole cell lysates. BRD4L-WT or BRD4 bromodomain mutant (BRD4L^mut^) Tet-On cells were treated with doxycycline 100 ng/ml overnight prior to dBET6 treatment (500 nM) for 2 hours. **i**, Effect of different BET bromodomain mutants on ER-mediated gene transcription measured by qPCR. Tet-On 3G:BET-BD cells were treated with doxycycline 10 ng/ml overnight prior to dBET6 treatment for 2 hours followed by 1-hour induction by E_2_. Data are shown as mean ± s.d. from independent triplicates.

**Extended Data Fig. 2.**
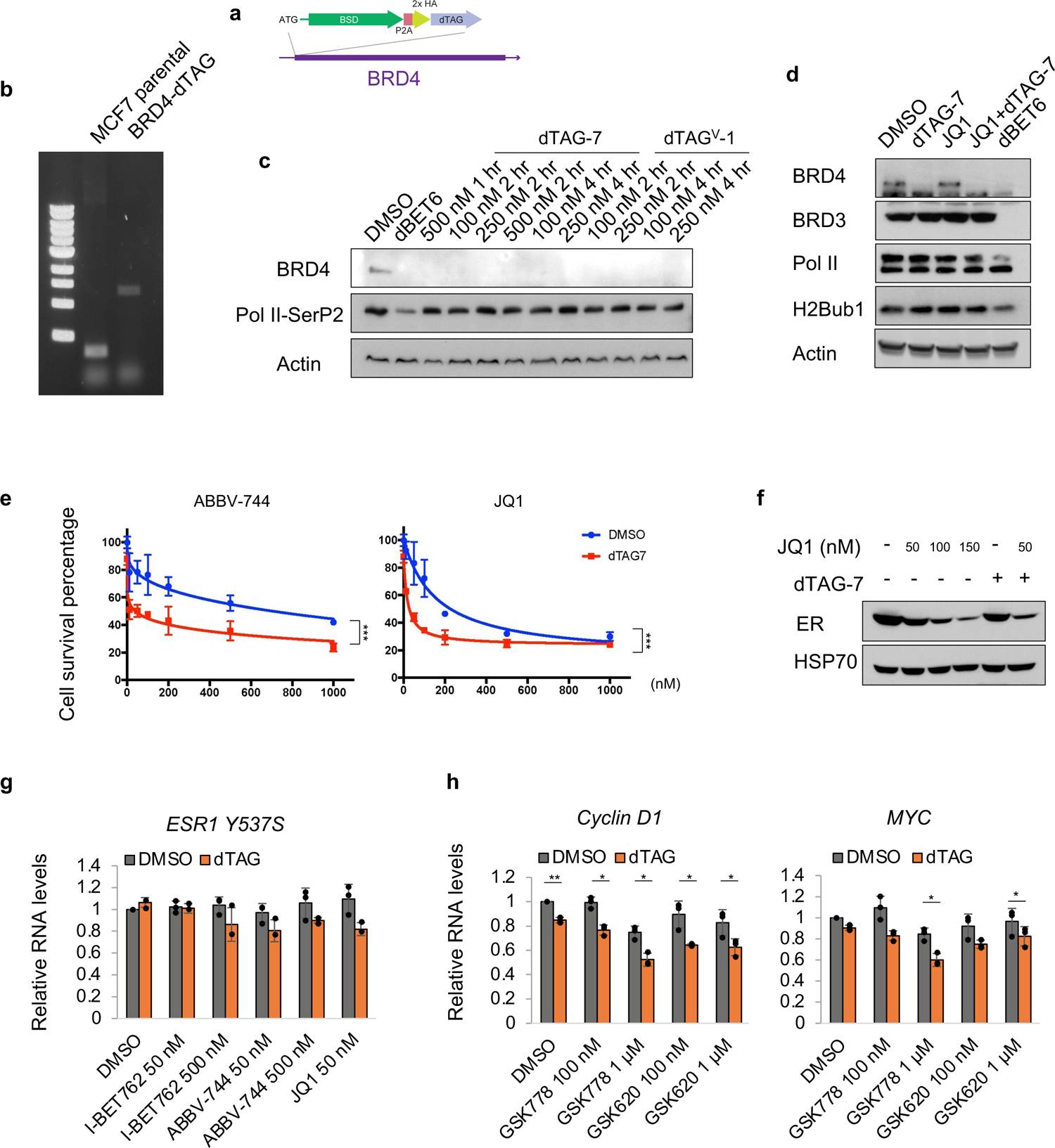
BRD4 mediates ER-dependent transcriptional resistance to bromodomain inhibition, related to Fig. 2. **a**, Schematic of the knockin strategy for BRD4-dTAG. **b**, Agarose gel electrophoretic analysis of PCR products from MCF7 parental cells and MCF7 BRD4-dTAG knockin cells. **c**, Immunoblot for BRD4 expression in BRD4-dTAG cells treated with dTAG-7 or dTAG^V^-1 at indicated concentrations for indicated times. dBET6 was added for 2 hours to show changes in Pol II CTD Ser2-P. **d**, Immunoblot showing effects of different 2-hour treatments on Pol II and H2Bub1 in BRD4-dTAG cells. **e**, Percentage of MCF7 BRD4-dTAG cell survival after treatment with or without BET inhibitors and dTAG-7 (250 nM) for 6 days. Data shown as mean ± s.d. from biological triplicates. **f**, Immunoblot showing reduced ER expression in BRD4-dTAG cells upon overnight treatment with JQ1 and dTAG-7. **g,h,** qPCR of *MYC, Cyclin D1* (**h**) and *ESR1-Y537* g, mRNA levels in MCF7 BRD4-dTAG ESR1-Y537S cells upon overnight treatment with BET inhibitors and dTAG-7 (500 nM). Because ESR1-Y537S-IRES-GFP is transcribed as a single transcript, *ESR1*-Y537S RNA level was monitored by GFP RNA. All cells were grown in normal full medium containing 10% FBS. Data shown as mean ± s.d. from independent triplicates. p values from two-tailed t-tests are depicted with asterisks.

**Extended Data Fig. 3.**
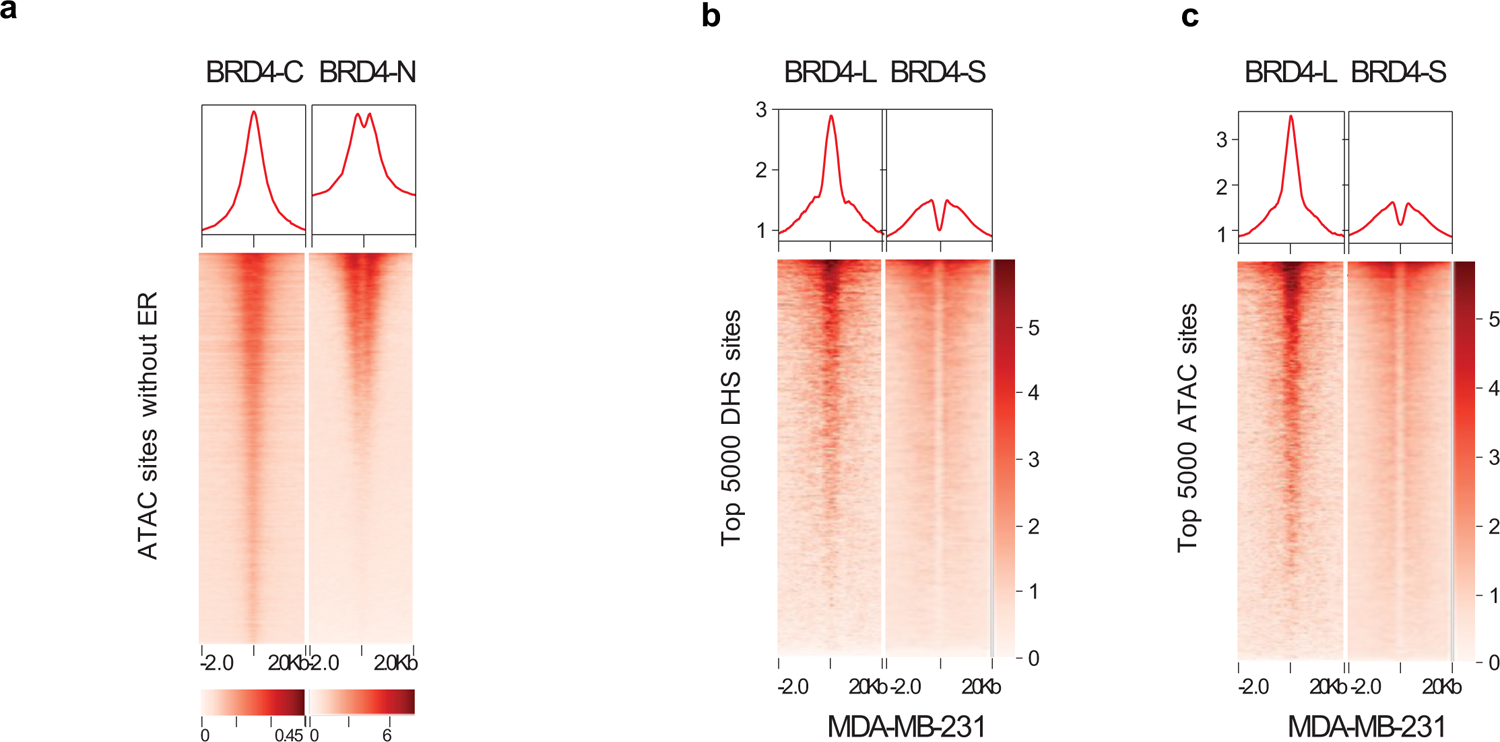
BRD4 isoforms occupy different genomic regions, related to Fig. 3. **a**, Heatmap of BRD4-N or BRD4-C ChIP-seq signals at non-ERBS ATAC sites in MCF7 cells. **b,c,** Analysis of public datasets showing that BRD4L is primarily recruited to nucleosome-free regions occupied by transcription factors, and that BRD4S tends to bind surrounding sites with acetylated histones. Density of ChIP-seq signals at the top 5,000 DNase hypersensitivity sites (DHS) (**b**) and the top 5,000 ATAC sites (**c**) in MDA-MB-231 cells. Endogenous BRD4-L was immunoprecipitated with a home-made antibody from the laboratory of Dr. Cheng-Ming Chiang (University of Texas, Southwestern). Overexpressed FLAG-tagged BRD4-S was immunoprecipitated with anti-FLAG^59^.

**Extended Data Fig. 4.**
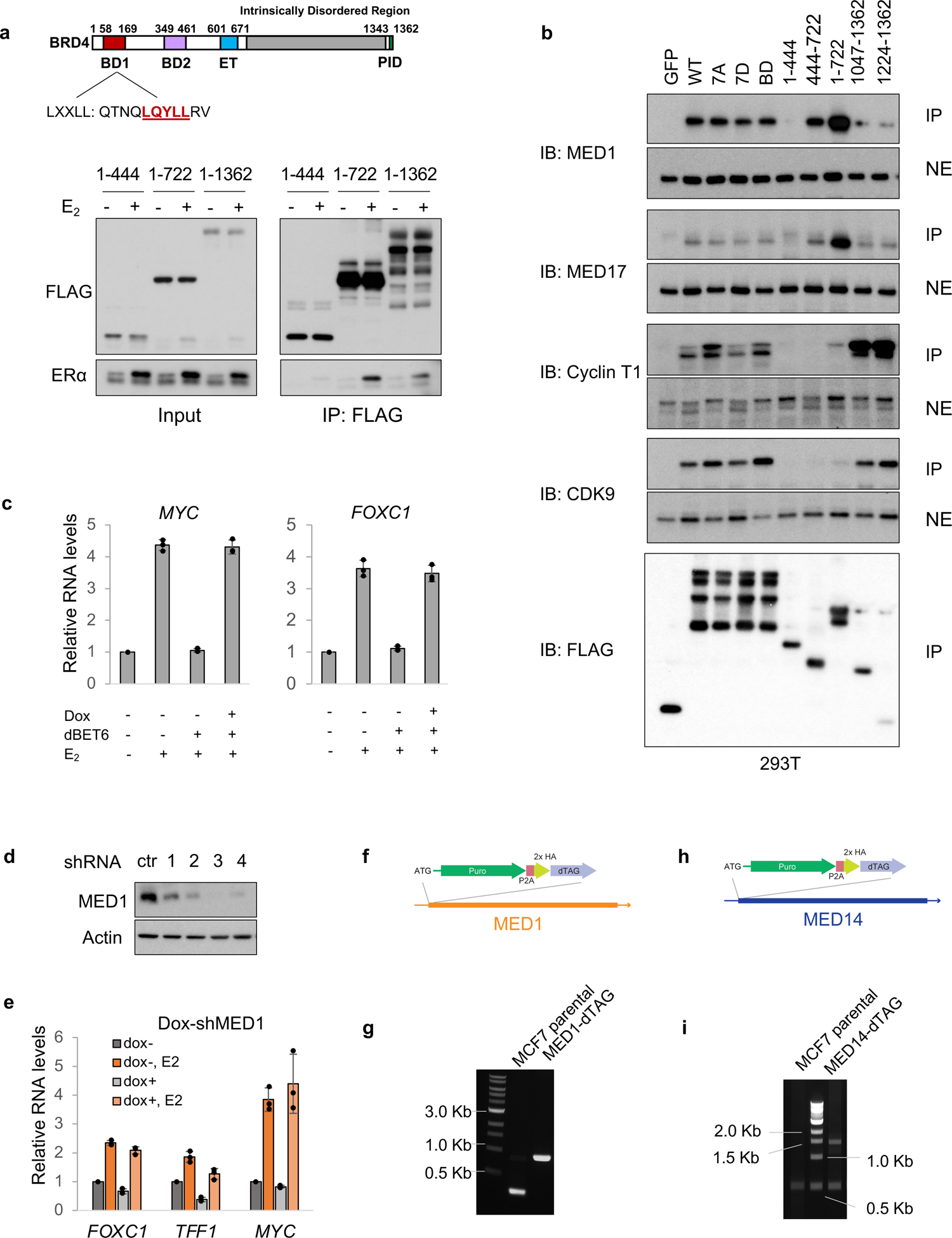
BRD4 associates with ER and Mediator, related to Fig. 4. **a**, Schematic of BRD4 indicating the two bromodomains (BD1 and BD2), the Extra-Terminal domain (ET), and the P-TEFb-interacting domain (PID) (upper panel). Immunoblot showing bromodomain- and LX motif-independent association of ER (lower panel). For this analysis, MCF7 cells in charcoal-stripped medium were transfected with vectors expressing FLAG-tagged BRD4 fragments (indicated at the top). After 2 days, E_2_ was added for 1 hour and nuclear extracts were collected and subjected to immunoprecipitation with FLAG antibody. **b**, Immunoblot of anti-FLAG immunoprecipitates of nuclear extracts from 293T cells transfected with FLAG-tagged GFP, BRD4, BRD4 mutants and BRD4 fragments (indicated at the top). **c**, Effect of BRD4 BD/ET double mutant (designated BRD4-BE) expression on ER-mediated gene transcription measured by qPCR. The BRD4-BE mutant was generated by mutating the ET domain (E651A/E653A) in the BRD4L^mut^ construct. The BRD4-BE mutant was incorporated into MCF7/FRT/Tet-on cells by FLP recombinase. The resulting BRD4-BE cells were treated the same as the BRD4L^mut^ cells in Fig. 1d. Cells were treated with doxycycline overnight to induce BRD4-BE expression and, subsequently, with dBET6 to deplete endogenous BET proteins. Cells then were treated with E_2_ for 1 hour to induce ER-dependent transcription. **d**, Immunoblot of MED1 protein levels in MCF7 dox-shMED1 cells. shRNA-3 was used for subsequent qPCR analysis. **e**, qPCR of RNA levels after MED1 knockdown indicating that MED1 is not required for ER-mediated transcription. Doxycycline (100ng/ml) was added to dox-shMED1 cells for 3 days prior to E_2_ stimulation for 1 hour. **f,** Schematic of the knockin strategy for MED1-dTAG. **g**, Agarose gel electrophoretic analysis of PCR products from MCF7 parental cells and MED1-dTAG knockin cells. **h**, Schematic of the knockin strategy for MED14-dTAG. **i**, Genomic PCR of a region spanning the start codon of *MED14* in MCF7 parental cells and MED14-dTAG cells. Data shown as mean ± s.d. from independent triplicates.

**Extended Data Fig. 5.**
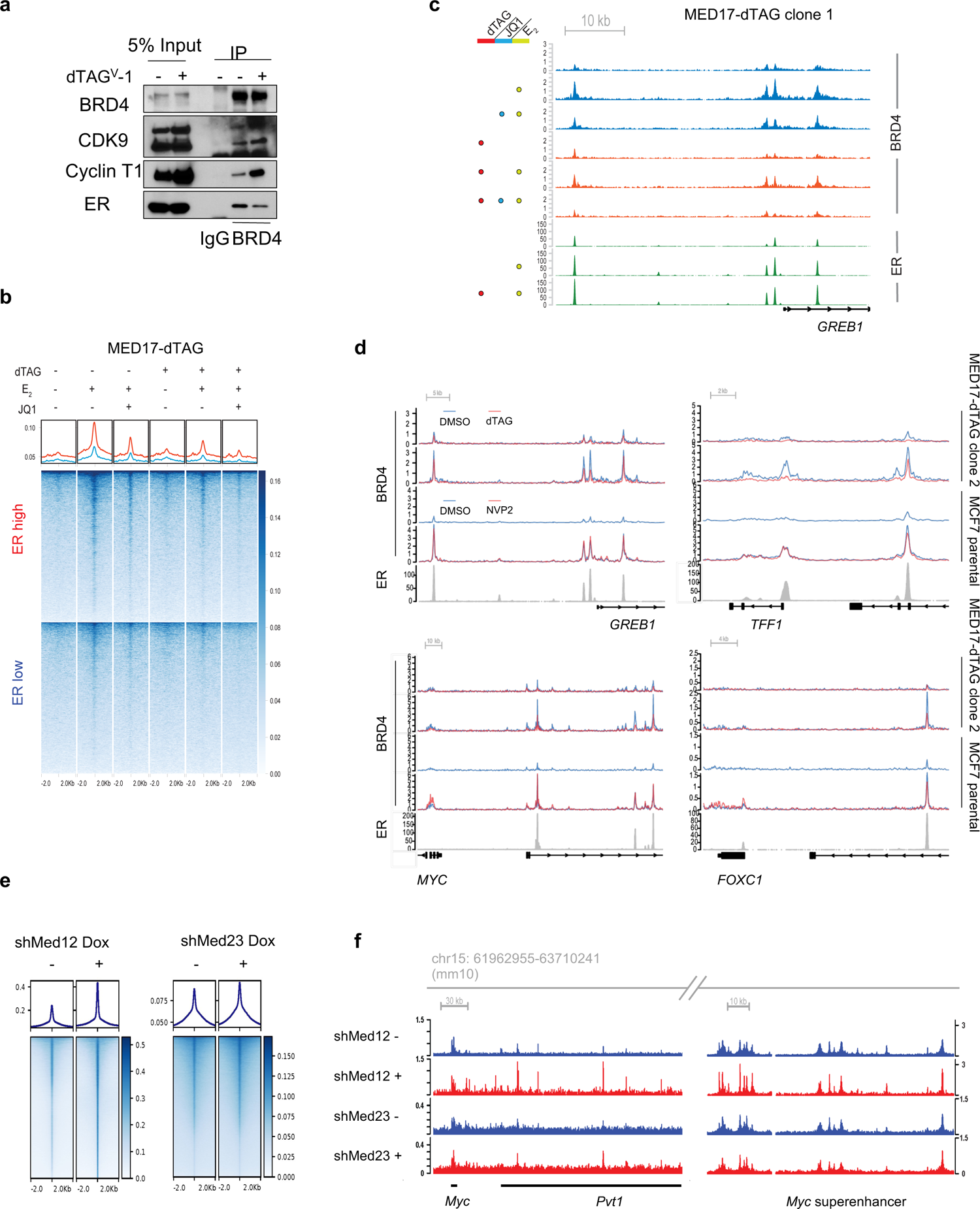
Mediator disruption affects BRD4 enhancer binding, related to Fig. 5. **a**, Immunoblot of indicated proteins from co-IP of BRD4 in nuclear extracts from MED17-dTAG cells treated with or without dTAG^V^-1 for 2 hours. Cells were grown in normal full medium. **b**, Heatmap of BRD4 ChIP-seq signals at ERBSs from MED17-dTAG clone 1 without DSG crosslinking. Cells were treated with or without 500 nM JQ1 or 500 nM dTAG^V^-1 prior to estrogen treatment. **c**, ChIP-seq signals of ER and BRD4 at the *GREB1* enhancer from MED17-dTAG clone 1 with DSG crosslinking. **d**, ChIP-seq signals of ER and BRD4 at different enhancers. **e,f,** Analysis of public dataset under accession number GSE78221 showing that knockdown of specific Mediator subunits increases Brd4 occupancy in MLL-AF9 transformed acute myeloid leukemia cells (RN2). ChIP-seq signals of Brd4 binding genome-wide (**e**) and at the *Myc* superenhancer (**f**).

**Extended Data Fig. 6.**
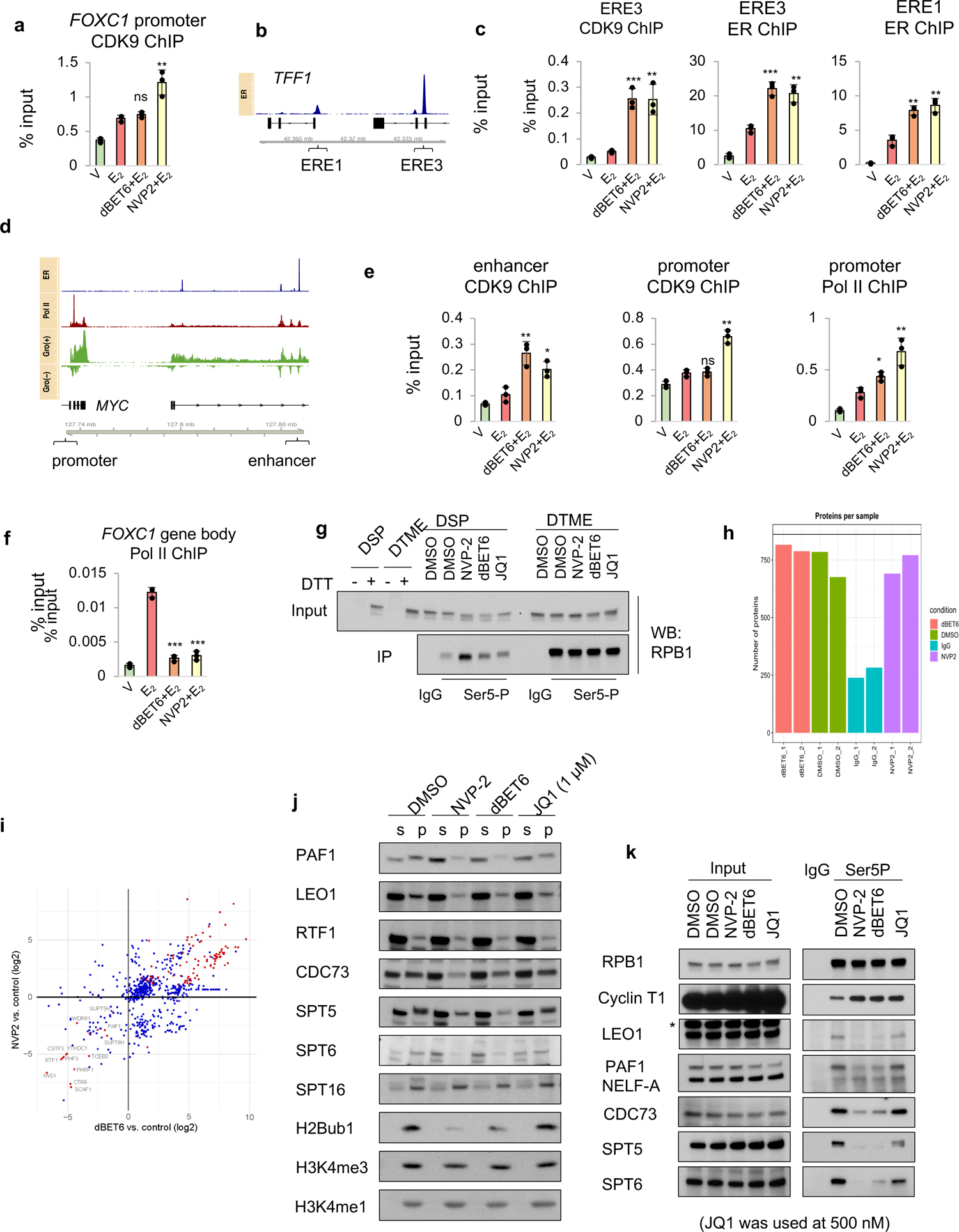
BET proteins depletion reduces elongation factor association with Pol II except P-TEFb, related to Fig. 6. **a**, ChIP-qPCR analyses of CDK9 binding at the *FOXC1* promoter. **b**, Gene track showing ER binding at enhancers of *TFF1*. **c**, ChIP-qPCR analyses of ER and CDK9 binding at the *TFF1* enhancers. **d**, Gene track showing ER and Pol II binding at the *MYC* enhancers and promoter. Gro-seq signals are also shown. **e**, ChIP-qPCR analyses of Pol II and CDK9 binding at the *MYC* promoter and a major enhancer. **f**, ChIP-qPCR of Pol II at the *FOXC1* gene body. Pol II ChIP was performed by using an RPB1 NTD antibody. For all ChIP analyses here, inhibitors or degraders were added 2 hours prior to E_2_ treatment for 40 min and are indicated in the figures. Data shown as mean ± s.d. from independent triplicates. p values from two-tailed t-tests (compared to E_2_ treatment alone) are depicted with asterisks. **g**, Immunoblot of Pol II showing (1) equal enrichment of Pol II CTD Ser5-P by DTME crosslinking but not DSP crosslinking under different treatment conditions and (2) higher enrichment efficiency by DTME relative to DSP. MCF7 cells treated with indicated inhibitors or degrader for 2 hours were crosslinked with DSP or DTME. After crosslinking, cells were collected and sonicated. Cell lysates were cleared by centrifugation and subjected to immunoprecipitation with Pol II CTD Ser5P antibody followed by immunoblotting analysis. **h**, Bar chart showing the number of proteins identified by the MaxQuant program in each sample from two independent replicates. **i,** Relative enrichment of proteins in dBET6 or NVP2 treatment compared with DMSO was calculated by the protein abundance from Proteome Discoverer. Protein abundance was normalized by RPB1. Data points with log2 fold change more than 1 in both conditions are highlighted in red. **j**, Immunoblots of indicated proteins confirming the loss of elongation proteins from the chromatin of cells treated with dBET6, JQ1 (1000 nM) or NVP2 for 2 hours. s, soluble fraction; p, insoluble pellet. The distribution of SPT16, a FACT component that facilitates elongation, was not affected by any treatment. **k,** Immunoblots of indicated proteins showing less change of elongation factors in JQ1 (500 nM) treatment than in dBET6 treatment. This is the same experiment as shown in (**g**) using DTME. Note that the crosslinking efficiency was very low for NELF-A and the signals in Ser5P IP were barely detected. The asterisk indicates an unknown band.

**Extended Data Fig. 7.**
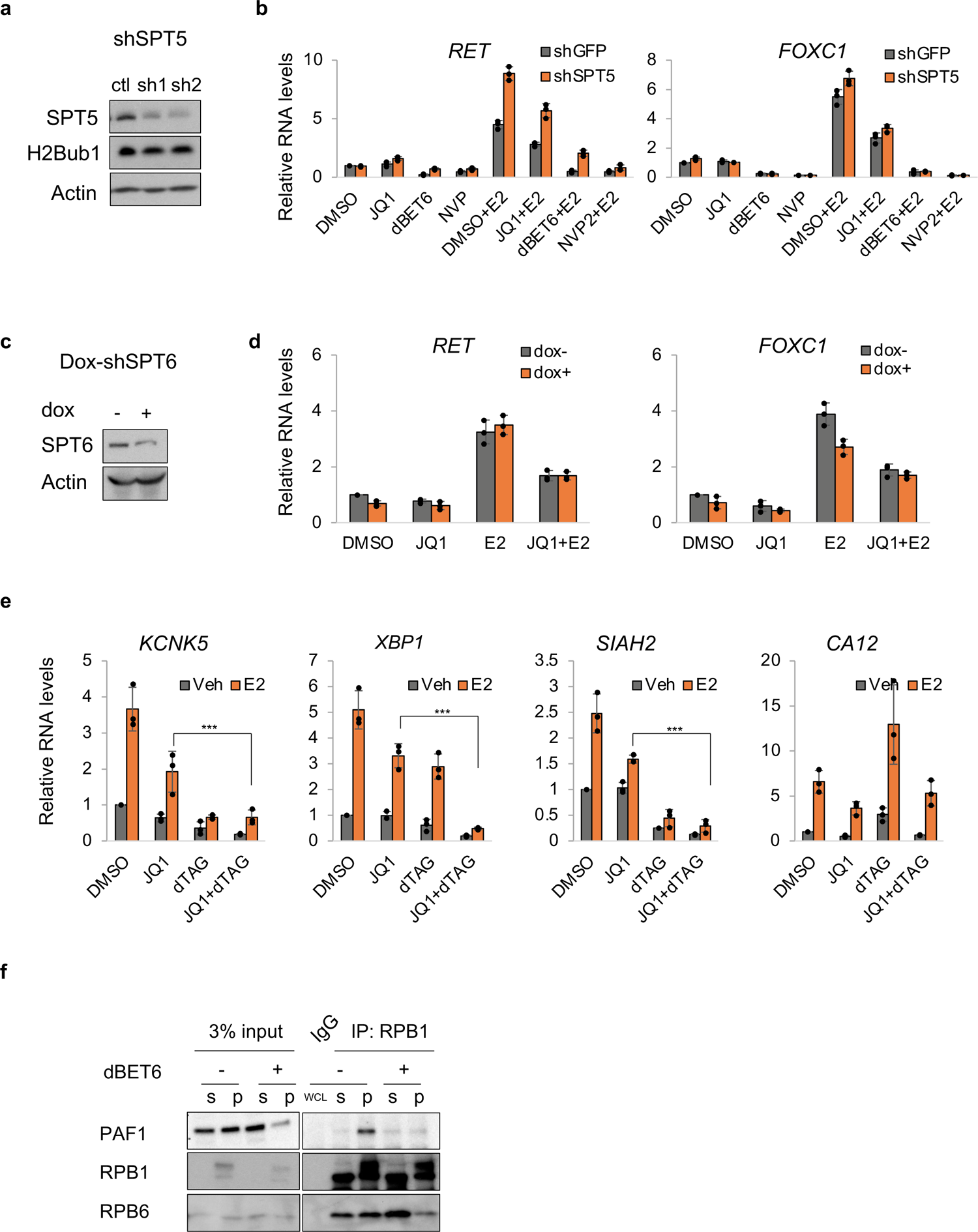
PAF1 mediates BETi-resistant ER-dependent transcription, related to Fig. 7. **a**, Immunoblot of SPT5 confirming SPT5 knockdown efficiency in MCF7 cells. **b**, qPCR of *FOXC1* and *RET* RNA levels in MCF7 cells expressing shRNAs targeting GFP or SPT5 and treated as indicated. Cells expressing SPT5-sh1 and SPT5-sh2 were pooled for RNA analysis. **c**, Immunoblot of SPT6 confirming SPT6 knockdown efficiency in MCF7 dox-shSPT6 cells treated with doxycycline for three days. **d**, qPCR of *FOXC1* and *RET* RNA levels in dox-shSPT6 cells with or without doxycycline treatment for three days and then treated as indicated. **e**, qPCR of gene expression changes in FLAG-PAF1-dTAG cells treated with or without JQ1 or dTAG^V^-1 for 2 hours prior to E_2_ stimulation. **f**, Immunoblot of indicated proteins from a Pol II co-IP using RPB1 NTD antibody showing reduced PAF1 association with Pol II in MCF7 cells following BET protein depletion by dBET6. Cells treated with or without dBET6 for 2 hours were subjected to fractionation into soluble fraction (s) and chromatin pellet (p). All fractions were then briefly sonicated and cleared by centrifugation followed by immunoprecipitation. WCL, whole cell lysates. Data shown as mean ± s.d. from independent triplicates. p values from two-tailed t-tests are depicted with asterisks.

## Notes

### Competing Interest Statement

The authors have declared no competing interest.

